# fMRI and MEG Fingerprints Diverge at the Individual Level

**DOI:** 10.64898/2026.04.27.721106

**Authors:** Brian Zhaoyi Mo, Stephen Smith, Mark Woolrich

## Abstract

Functional connectivity (FC) profiles derived from fMRI and MEG offer complementary perspectives on large-scale brain organization, while showing reasonable correspondence at the population-average level. However, how their individual variability relates between these modalities remains unclear. Using the Cam-CAN dataset, we derived neural fingerprints from subject-level resting-state fMRI FC and MEG FC obtained from the same participants (N=543). Fingerprints derived from each modality separately showed robust within-subject, cross-session consistency and successfully predicted age and cognition, confirming that these features capture stable and behaviourally relevant individual traits. We then quantified shared individual variability between modalities using variance partitioning analyses and representational similarity measures. Two main findings emerged. First, despite strong similarity at the population-average level, correspondence between MEG and fMRI neural fingerprints at the subject level was low, as reflected in both cross-modal shared variance and the preservation of pairwise inter-subject similarity patterns, quantified by linear Centred Kernel Alignment (CKA). Second, structural fingerprints accounted for the majority of age-related variance in functional neural fingerprints, almost entirely explaining the age-related variance in, and shared between, fMRI and MEG. MEG functional fingerprints did have unique information not accounted for by structure when explaining variability in cognitive traits, but this was not shared with fMRI. Together, these findings demonstrate that there is a surprisingly lack of similarity in the way that subjects vary between fMRI and electrophysiology, especially when structural variability is accounted for.

## 1. Introduction

The human brain exhibits rich intrinsic activity, organized into large-scale functional networks that shape cognition and behaviour. Characterizing these networks is central to understanding individual differences in healthy aging and disease. Functional magnetic resonance imaging (fMRI) and non-invasive electrophysiology (e.g., magneto-/electroencephalography, M/EEG) have served as primary windows into this organization. fMRI detects fluctuations in blood-oxygen-level-dependent (BOLD) response, thereby revealing large-scale distributed networks in spontaneous brain activity. While fMRI reveals large-scale distributed networks by capturing slow, hemodynamically coupled blood-oxygen-level-dependent (BOLD) fluctuations with high spatial precision (Biswal et al., 1995; Fox & Raichle, 2007; Shulman et al., 1997), MEG directly measures electrophysiological oscillations with millisecond temporal resolution (De Pasquale et al., 2010; Mantini et al., 2007). Despite these differences, both modalities have independently revealed that individual functional brain organization is unique and stable over time, a phenomenon known as “neural fingerprinting” (Finn et al., 2015; da Silva Castanheira et al., 2021).

At the group level, fMRI and MEG reveal similar functional brain network architecture. Time-averaged connectivity patterns exhibit a notable degree of similarity between M/EEG and fMRI when analysing MEG oscillatory power in specific frequency bands, particularly in the alpha and beta bands (Brookes et al., 2011; Hall et al., 2014). Dynamic modelling of M/EEG data further reveals transient resting-state networks (RSNs) that are consistent at the group level with those identified via fMRI (Baker et al., 2014; Cho et al., 2024).

Despite this strong group-level convergence, it remains unclear whether such correspondence holds at the level of the individual. Does the individual with the strongest connectivity in the hemodynamic domain also exhibit the strongest connectivity in the electrophysiological domain? Establishing this cross-modal correspondence is critical: if hemodynamic and electrophysiological fingerprints align, they likely reflect a unitary source of neural variability; if they diverge, they offer complementary, non-redundant windows into individual brain function.

Furthermore, any observed relationship between fMRI and MEG variability must be disentangled from underlying anatomy. Structural features such as cortical thickness and white matter integrity constrain both hemodynamic and electrophysiological activity and are themselves predictive of age, behaviour, and other demographic variables (Llera et al., 2019), potentially driving the relationship between both functional modalities. It therefore remains an open question whether any convergence between fMRI and MEG fingerprints reflects a shared latent neural source that is expressed in both hemodynamic and electrophysiological signals, or whether it instead arises from anatomical properties that influence both signals in parallel.

A particular concern is that individual differences in the size, shape, and exact cortical position of brain regions interact strongly with the modelling of functional connectivity, such that cross-subject variability in spatial topography can be interpreted as variability in FC edges rather than reflecting genuine differences in neural coupling (Bijsterbosch et al., 2018). Unless such anatomical structural sources are accounted for, the correspondence between fMRI and MEG fingerprints may be overestimated, particularly in the context of ageing.

To address these questions, we leveraged the Cambridge Centre for Ageing and Neuroscience (Cam-CAN) dataset (N=612) to systematically map the relationship among structural, hemodynamic and electrophysiological variability. First, we derived FC fingerprints from resting-state fMRI and MEG data and established the validity of these metrics by demonstrating that fingerprints derived from both modalities are robust, reliable, and predictive of age and fluid intelligence. Second, we demonstrate that, despite the similarity of fMRI and MEG functional networks at the group level, fMRI and MEG fingerprints exhibit surprisingly limited direct correspondence at the individual-subject level. Third, using variance partitioning (Ray-Mukherjee et al., 2014; Seibold & McPHEE, 1979), we tested the hypothesis that cross-modal convergence is heavily driven by anatomy. Our results show that structural factors explain most of the shared age-related variance between fMRI and MEG; however, they do not account for the total variability, leaving a substantial portion of variance unique to each functional modality.

Taken together, our results show that hemodynamic and electrophysiological profiles capture largely distinct sources of individual variability, necessitating a multimodal approach to fully characterize brain functional connectivity.

## 2. Methods

### 2.1 Dataset

We utilized the Cambridge Centre for Ageing and Neuroscience (Cam-CAN) dataset in this project, in which both resting-state fMRI, MEG and T1w MRI are available in the same subjects among the healthy cohort (N=612; age range: 18-88 years)(Shafto et al., 2014; Taylor et al., 2017). After image processing, there are 543 subjects available that have data from all modalities. To facilitate cross-modality comparative analysis, we applied the Glasser52 parcellation (Kohl et al., 2023), consisting of 52 parcels, to both modalities. The Glasser52 parcellation was derived from Human Connectome Project Multimodal Parcellation (HCP-MMP) by estimating the contribution of each region to the MEG sensor signal and then merging and subdividing regions so that the 52 results parcels contribute comparable magnitude to the MEG signal. This reduces dimensionality to a level appropriate for MEG while balancing signal-to-noise across parcels. The Glasser52 parcellation provides a common space, in which both modalities can be summarised at matched parcels, and therefore supports direct cross-modal comparisons at the parcel level. For fMRI, we also adopted HCP-style surface processing, one of the most widely used frameworks for fMRI analysis. In addition to the Glasser52 parcellation, individual-level fMRI data were also parcellated in surface space using ICA dual regression based on the HCP group-ICA maps with 25-and 50-component decompositions. The motivation for this second representation is that fMRI and MEG express individual variability most strongly under different processing pipelines; the surface space representations therefore allows fMRI to contribute its most informative features to the cross-modal comparisons.

### 2.2 Data acquisition and preprocessing

#### 2.2.1 structural MRI data acquire and processing

The structural MRI data were acquired using a 3D T1-weighted MPRAGE sequence (TR = 2250 ms, TE = 2.99 ms, TI = 900 ms, flip angle = 9°, FOV = 256 × 240 × 192 mm, 1 mm isotropic resolution, GRAPPA = 2, acquisition time = 4 min 32 s).

T1-weighted images were processed using an adapted UK Biobank (UKB) pipeline (Alfaro-Almagro et al., 2018), which performs gradient distortion correction, field-of-view cropping, brain extraction, defacing, and registration (linear and non-linear) to MNI152 space, followed by tissue segmentation. The processed T1-weighted image served as the anatomical reference for co-registering all other imaging modalities.

From the T1 images, we derived imaging-derived phenotypes (IDPs) including global and regional volumetric measures (total brain, grey matter, white matter, ventricular CSF, and peripheral cortical grey matter volumes), as well as 139 regional grey matter volumes and bilateral subcortical volumes (e.g., thalamus, caudate, hippocampus). Additional morphometric IDPs, such as cortical thickness, area, and volume, were obtained via FreeSurfer (Fischl, 2012) using three structural atlases.

#### 2.2.2 fMRI data acquisition and processing

The resting-state fMRI data were acquired using a T2\***-**weighted gradient-echo EPI sequence (261 volumes, 32 axial slices, slice thickness = 3.7 mm, interslice gap = 0.74 mm, TR = 1970 ms, TE = 30 ms, flip angle = 78°, FOV = 192 × 192 mm², voxel size = 3 × 3 × 4.44 mm, total acquisition time = 8 min 40 s).

Preprocessing followed the Cam-CAN adapted UKB functional pipeline (Alfaro-Almagro et al., 2018), incorporating several key steps: unwarping to correct magnetic field inhomogeneities, gradient distortion correction (GDC), motion correction using MC-FLIRT (Jenkinson et al., 2002), and slice-timing correction. Functional images were then co-registered to each participant’s T1-weighted structural image, normalized to MNI152 space, and spatially smoothed using a 5 mm FWHM Gaussian kernel.

Artifact removal was performed using ICA+FIX (Beckmann & Smith, 2004; Griffanti et al., 2014; Salimi-Khorshidi et al., 2014). A Cam-CAN-specific FIX classifier was hand-trained on 20 randomly selected resting-state and task fMRI datasets following (Griffanti et al., 2017). Leave-one-out validation yielded mean accuracies of 100% for non-artifact and 95% for artifact components, confirming robust noise classification.

In volumetric space, denoised fMRI data were regressed onto parcellation spatial maps to extract subject-specific time series for each parcel. In surface space, data underwent HCP-style surface-based processing, followed by ICA dual regression using HCP group-ICA maps with 25-and 50-component decompositions to derive subject-specific timeseries.

#### 2.2.3 MEG data acquisition and processing

The MEG raw data were collected using a 306-channel Elekta Neuromag Vectorview system, which includes 102 magnetometers and 204 planar gradiometers. The data were sampled at 1 kHz with a highpass filter of 0.03 Hz. The scanning time of MEG was at least 8 minutes and 40 seconds, to match the fMRI scanning time.

The MEG preprocessing pipeline was implemented in osl-ephys (Quinn et al., 2022; Van Es et al., 2025) and involved four steps: (1) Filtering: bandpass filtering of 0.5-125 Hz, notch filtering at 50 and 100 Hz to remove powerline noise, and notch filtering at 88 Hz to remove spike artefact; (2) Downsampling: the data were downsampled to 250 Hz; (3) Automated bad segments and channel detection; (4) Artefact Removing: Independent Component Analysis (ICA) with 64 components was used to identify artefacts.

The MEG data was first co-registered with individual T1 structural images, to align MEG sensor positions to the cortical surface. The preprocessed sensor data were bandpass filtered between 1-45 Hz and then source reconstructed onto an 8mm isotropic grid using a linearly constrained minimum variance (LCMV) beamformer (Van Veen & Buckley, 1988). Finally, Parcel-wise signals were derived by applying principal component analysis (PCA) to the source space data, with each parcel’s time series represented by the first principal component, capturing the majority of variance across its voxels. Symmetric orthogonalization was applied to prevent spatial leakage.

### 2.3 Time Delayed Embedded Hidden Markov Model

HMM is a model that can partition multi-channel data into a finite number of states. Each HMM state represent a dynamic network, which has a distinct spatial-temporal pattern. The generative model of HMM includes two parts: a temporal model which includes a time course indicating which of the hidden states is active, and a spatial observation model for each state (Gohil et al., 2024). In the temporal model, using a Markov process, states time courses are derived from a transition probability matrix. In the observation model for each state, in this work we assume that the data are generated by a multivariate Gaussian distribution, in which the means and covariances describe the within and between channel characteristics for each state.

The inference of HMM parameters was carried out using a variant of the Baum-Welch algorithm, which is an application of the Expectation-Maximization algorithm (EM) for HMM. In the “Expectation” step, the states’ probabilities were updated. In the “Maximization” step, with the updated state probabilities, the transition probability matrix were updated with a stochastic updates technique, and other parameters were updated by minimizing the variational free energy (Gohil et al., 2024).

Consistent with previous work, the channel data we fed into the HMM for inference corresponds to the parcel MEG time-courses (typically after they have been corrected for spatial leakage). However, we augmented the parcel time courses with time delay embedding (TDE) using ± 7 lags to allow the HMM to find states characterised by dynamics in oscillatory amplitude and phase synchronization between channels (Gohil et al., 2024). With a 52-ROI parcellation (Glasser52), this embedding resulted in a total of 780 channels. PCA was then applied to the TDE parcel data to reduce the dimensionality down to 100 channels. The data were also standardised so that each channel’s signal had a zero mean and variance of one.

The TDE-HMM was trained with 8 states for each session. Input sequences of 400 samples were processed in batches of size 256. Then we used Adam optimizer to learn the trainable parameters with learning rate of 0.01 over 25 epochs. The HMM model was trained 5 times for each session, and the optimal one with lowest free energy among these 5 runs was selected.

We then obtained estimates of subject-specific HMM state means and covariances using dual estimation (Gohil et al., 2024; Huang et al., 2026). Next, we backprojected the PCA we had carried out on the TDE parcel data, to get the means and covariances in the original TDE channels space.

In this work, we ran separate HMMs on the different subsets (i.e., sessions) of the data. Each HMM therefore had arbitrary state orderings. To allow comparisons between the HMMs from different sessions, we reordered (i.e., aligned) the states using cross-session state-to-state covariances correlation and the Hungarian algorithm to determine the optimal pairings based on the correlation coefficients.

### 2.4 Network features analysis

We extracted a total of 23 functional features (Table 1) from fMRI and MEG.

**Table 1:** All extracted features:

| Abbreviation | Feature type |
| --- | --- |
| fmri_surface_d50_full | fMRI surface space (50 PCs, full correlation) |
| fmri_surface_d50_partial | fMRI surface space (50 PCs, partial correlation) |
| fmri_surface_d25_full | fMRI surface space (25 PCs, full correlation) |
| fmri_surface_d25_partial | fMRI surface space (25 PCs, partial correlation) |
| fmri_parcel_full | fMRI volumetric space Glasser52 (full correlation) |
| fmri_parcel_partial | fMRI volumetric space Glasser52 (partial correlation) |
| meg_aec_broad | MEG amplitude envelope correlation (broadband, 1-45 Hz) |
| meg_aec_delta | MEG source space, amplitude envelope correlation (delta band, 1-4 Hz) |
| meg_aec_theta | MEG source space, amplitude envelope correlation (theta band, 4-8 Hz) |
| meg_aec_alpha | MEG source space, amplitude envelope correlation (alpha band, 8-13 Hz) |
| meg_aec_beta | MEG source space, amplitude envelope correlation (beta band, 13-24 Hz) |
| meg_aec_gamma | MEG source space, amplitude envelope correlation (gamma band, 30-45 Hz) |
| meg_parcel_tde_cov | MEG source space, time-delay embedded covariance |
| meg_hmm_covs | MEG source space, HMM state covariances (dynamic) |
| meg_sensor_cov | MEG sensor space, covariance |
| meg_sensor_tde_cov | MEG sensor space, time-delay embedded covariance |
| meg_nc_aec_broad | No leakage correction MEG amplitude envelope correlation (broadband) |
| meg_nc_aec_delta | No leakage correction MEG amplitude envelope correlation (delta band) |
| meg_nc_aec_theta | No leakage correction MEG amplitude envelope correlation (theta band) |
| meg_nc_aec_alpha | No leakage correction MEG amplitude envelope correlation (alpha band) |
| meg_nc_aec_beta | No leakage correction MEG amplitude envelope correlation (beta band) |
| meg_nc_aec_gamma | No leakage correction MEG amplitude envelope correlation (gamma band) |
| meg_nc_parcel_tde_cov | No leakage correction MEG parcel-level time-delay embedded covariance |
| struct | Structural IDPs |
| struct_FS | FreeSurfer-derived structural IDPs (thickness and area) |
| struct_concat | Concatenated structural feature set |

#### 2.4.1 fMRI FC

For fMRI, we estimated fMRI FC using two methods for three different parcellation setups:

1. Time-averaged (i.e., static) full FC, computed as the full correlation matrix, then a fisher z-transformation was used to convert correlation to approximately normal distribution. The group level FC was calculated as the average across subjects.
2. Time-averaged (static) partial FC, computed as the partial correlation matrix, then a fisher z-transformation was used to convert correlation to approximately normal distribution. The partial correlated was estimated using Tikhonov-regularized partial correlation, derived from the inverse of the covariance matrix with a ridge regularization term to improve numerical stability and mitigate noise amplification. The optimal regularization coefficient was determined by maximizing the Pearson’s correlation coefficient between each subject’s regularized precision matrix and the group-average of the unregularized precision matrices. The group level FC was calculated as the average across subjects.

#### 2.4.2 MEG FC

For MEG, we estimated MEG FC using four different methods:

1. Time-averaged (static) FC, obtained using Amplitude Envelope Correlation (AEC) across broad-band frequencies (1-45 Hz) and specific frequency bands: delta (1-4 Hz), theta (4-8 Hz), alpha (8-13 Hz), beta (13-24 Hz), and gamma (30-45 Hz). AEC can provide robust resting state MEG functional connectivity, and shows high test-retest reliability (Colclough et al., 2016). The static FC were also obtained as the TDE covariances with ± 7 lags. With a 52-ROI parcellation, this embedding resulted in a total of 780 channels. PCA was then applied to reduce the dimensionality down to 100 channels before standardization. We then calculated the covariance matrix of these channels.
2. Dynamic FC, obtained using the Time-Delay Embedded Hidden Markov Model (TDE-HMM) with dual estimation (Huang et al., 2026; Vidaurre et al., 2018);
3. Sensor space FC (static), obtained using covariances and TDE covariances; PCA was applied to reduce the sensor space data from 306 to 60 channels, which is approximately the rank of MEG sensor space data after MaxFiltering, to reduce memory overhead in the following computation. TDE-PCA covariances were also calculated in sensor space with the same setup as in source space with ± 7 lags and 100 components PCA reducing dimensionality.
4. No leakage corrected static FC, obtained in the same way as the static FC setup, but using source reconstructed data without applying symmetric orthogonalization.

### 2.5 Analysis on extracted features

#### 2.5.1 Group-level analysis

To examine the cross-modality FC’s properties, we first examined the similarities between the group level patterns. Where the fMRI group level FC and MEG group level FC used the same parcellation, the FCs between two modalities shared the same shape of 𝑛_𝑝𝑎𝑟𝑐𝑒𝑙𝑠_ × 𝑛_𝑝𝑎𝑟𝑐𝑒𝑙𝑠_ (52×52). Hence, we can examine their similarities by simply calculating the correlation between the off-diagonal of the FC matrices. Note that we cannot do this with the FC matrices extracted from TDE MEG data, as they do not share the same shape with fMRI FC matrices. In addition, we fitted a General Linear Model (GLM) between fMRI group FC patterns and MEG static specific bands’ FC patterns, to probe the relationship between fMRI and MEG group level patterns. We then correlated the GLM predicted FC matrix with fMRI group FC matrix.

#### 2.5.2 Individual-level analysis

All parameters for the individual-level analyses are outlined in Table 3, including inputs, output metrics, and the research questions addressed.

After computing the cross-modalities group-level similarity, we moved on to individual-level analysis. The FC for an individual subject can be modelled as being made up of: a) the group-mean FC for the modality in question, b) the within-modality, between-subject variability, and c) the within-subject, cross-session variability. To focus on subject-specific features, we first demeaned each subject’s FC by subtracting the group-average FC, yielding connectivity fingerprints that minimize shared variance with group-level structure and reflect clearer individual differences within each modality. Prior to subsequent analyses, the demeaned FC matrices were vectorized and reduced in dimensionality using principal component analysis (PCA), retaining the top 1,000 components to preserve the majority of variance while improving computational efficiency.

##### 2.5.2.1 Cross-session Robustness of Fingerprints

Within-subject, between-session variability reflects both measurement-related noise and genuine session-level fluctuation. The present design cannot separate these sources, and they are therefore treated together as nuisance variance in the subsequent analyses. To quantify the discriminability of fingerprints, we used cross-session subject label prediction. For each modality, we first calculated the subject similarity matrix, 𝐴, between sessions using Pearson correlation coefficients:

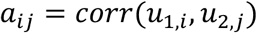

where 𝑢_𝑘,𝑖_ denotes subject 𝑖’s fingerprint pattern in session 𝑘. The diagonal area of 𝐴 represents subjects’ self-similarity between sessions, while the off-diagonal area is the between-subject similarity. The aim here is to predict the correct subject label in the target session (session 2), using the fingerprints in the source session (session 1). With the subject similarity matrix, 𝐴, we calculate the between-session subject identification accuracy as:

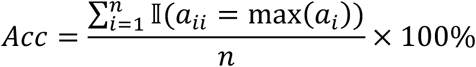

Where 𝑛 is the number of subjects. We evaluated the identification accuracy for each functional fingerprint’s features across two sessions.

We also calculated the linear Centred Kernel Alignment (CKA) between two sessions for each fingerprint. Cross-session subject identification and linear CKA probe two different levels of similarity. Identification is a first-order test: for each subject, it asks whether the fingerprint from session one is more similar to that same subject’s fingerprint in session two than to any other subjects, that is, whether two fingerprints themselves match across sessions. Linear CKA is a second-order test: it compares the two fingerprint spaces directly, asking whether the pairwise similarity among subjects in session one is preserved in session two.

Linear CKA therefore provides an estimate of the between-session consistency of the subject similarity matrix, as it is mathematically equivalent to the correlation between the centred similarity matrices derived from two sessions (see Supplementary Information for details).

The two metrics are complementary, and a feature may succeed on one and fail on the other. To calculate linear CKA, we first calculated the subject similarity matrix 𝐵 for each session:

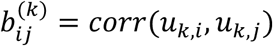

where 𝐵^(𝑘)^denotes the subject similarity matrix in session 𝑘, and 𝑢_𝑘,𝑖_ denotes subject 𝑖’s fingerprint pattern in session 𝑘. Notably, the diagonal elements of 𝐵^(𝑘)^ are 1s as it is a self-correlation matrix. We focused on the off-diagonal area of 𝐵^(𝑘)^ here, as it reflects the pairwise similarity patterns between subjects in each session, referred as the inter-subject similarity. Finally, by correlating the inter-subject similarity between sessions for each modality, we obtain the cross-session linear CKA, indicating how well the similarity for each fingerprint is preserved between sessions.

##### 2.5.2.2 Downstream prediction tasks using fingerprints

###### Age Prediction

To further interpret and compare the subject variability in the fingerprint, we used the extracted fingerprints to do age prediction in Cam-CAN, which has subjects with age ranging from 18 to 88 years. We applied age prediction with 10-fold cross-validated ElasticNet, with the regularisation strength and the L1-to L2 mixing ratio selected within each outer fold by nested 5-fold cross-validation on the training subjects only and evaluated using the coefficient of determination (𝑅^2^).

###### Cognition Prediction

To evaluate the behavioural relevance of each fingerprint, we performed prediction analyses on a set of cognitive tasks from Cam-CAN covering domains including face and emotion processing, memory, reasoning, executive function, and language (Table 2). Task scores were first deconfounded for age using linear regression, and the residuals were concatenated across tasks for all subjects with all relevant data present (n = 375). The combined scores were standardized and subjected to principal component analysis (PCA). The top four principal components (each explaining > 5% of variance) were retained as cognitive components.

**Table 2:** Description of cognitive tasks selected.

| Task | Domain | Brief Introduction |
| --- | --- | --- |
| BentonFaces | Face perception / recognition | Assesses face perception and recognition ability using unfamiliar and familiar face stimuli. |
| Cattell | Fluid intelligence / reasoning | Measures fluid intelligence through culture-fair reasoning tasks minimizing linguistic and educational bias. |
| EkmanEmHex | Emotion recognition (6 emotions) | Evaluates accuracy and response speed in recognizing six basic facial expressions of emotion. |
| FamousFaces | Famous face recognition / naming | Assesses semantic memory and name retrieval for well-known individuals from visual facial cues. |
| Hotel | Executive function | Examines multitasking and goal-management abilities under time-constrained real-life task demands. |
| PicturePriming | Implicit memory | Assesses implicit memory through response time facilitation for phonologically or semantically primed stimuli. |
| Proverbs | Abstract reasoning / language | Evaluates abstraction and verbal reasoning ability through interpretation of figurative proverbs. |
| RTchoice | Choice reaction time / attention | Measures motor speed and attentional control through rapid choice reactions to visual stimuli. |
| RTsimple | Simple reaction time | Assesses basic sensorimotor speed via reaction times to a single repeated stimulus. |
| Synsem | Sentence processing (syntactic/semantic ambiguity) | Examines syntactic and semantic processing efficiency in resolving linguistic ambiguities. |
| TOT(Tip-of-the-tongue) | Memory / name retrieval | Measures word-finding and lexical retrieval failures through frequency of tip-of-the-tongue states. |
| VSTMcolour | Visual short-term memory | Assesses capacity and precision of visual short-term memory using colour recall with variable set sizes. |

**Table 3:** Overview of individual-level analysis parameters: inputs, output metrics, and corresponding research questions, ordered according to the sequence of studies.

| Analysis | Input | Output/metric | Question addressed |
| --- | --- | --- | --- |
| Cross-session subject identification | Single modality, cross-session fingerprints | Identification accuracy | First-order test: does each subject's fingerprint match their own across sessions? |
| Cross-session similarity of inter-subject similarity | Single modality, cross-session fingerprints | Linear CKA | Second-order test: is the inter-subject similarity preserved across sessions? |
| Age prediction | Single modality fingerprints; age | Cross-validated $R^2$ | How much age-related variance does each fingerprint carry? |
| Cognition prediction | Single modality fingerprints; Cognition component scores | Cross-validated Pearson $r$ | Which fingerprints predict each cognitive component? |
| Age-related variance partitioning | Cross-modal triplet fingerprints' age-prediction $R^2$ | Estimated unique and shared $R^2$ sets | How is age-related variance distributed across modalities? |
| General variance partitioning | Cross-modal triplet fingerprints | Estimated unique and shared variance components | How much overall subject variability is shared across modalities? |
| Cross-modal shared variance | Cross-modal pairwise fingerprints | Percentage of variance explained | How much subject variability in target fingerprint can be explained by source fingerprints? |
| Cross-modal correspondence of inter-subject similarity | Cross-modal pairwise fingerprints | Linear CKA | Second-order test: is there any correspondence of inter-subject similarity between modalities? |
| Cross-modal subject identification | Cross-modal pairwise fingerprints | Identification accuracy | First-order test: how well can each subject's fingerprint match their own across modalities? |

Prediction was performed using 10-fold cross-validated ElasticNet regression, with the regularisation strength and the L1-to L2 mixing ratio selected within each outer fold by nested 5-fold cross-validation on the training subjects only, and accuracy was quantified by the correlation between predicted and observed cognitive component scores. Statistical significance was assessed using a permutation test with 1000 resamples, with maximum-statistics correction across components to achieve family-wise error control for multiple comparisons.

##### 2.5.2.3 Cross-modal fingerprints alignment

###### Age-related variance partitioning

We quantified the shared and unique age-related variance captured by the three modalities (A: fMRI, B: MEG, C: structural MRI) based on their age-prediction results. For each modality and their combinations, we obtained the coefficient of determination (𝑅^2^) between the predicted and chronological age using cross-validated predictions, calculated as:

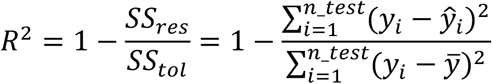

These 𝑅^2^ values were treated as sets representing the observed age-related variance explained by each modality.

Following the framework of set theory, the unique and shared variance components were computed as intersections and differences among these sets. For example:

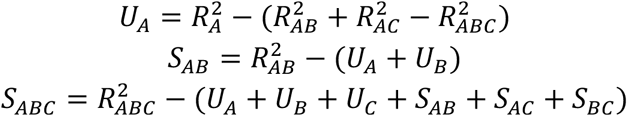

where 𝑈 denotes the estimated unique variance of each modality, 𝑆_𝐴𝐵_ the estimated variance shared between A and B only, and 𝑆_𝐴𝐵𝐶_ the estimated variance jointly shared by all three modalities.

All 𝑅^2^values were derived from cross-validated predictions. The resulting partitions were normalized by the total explained variance 𝑅^2^ and visualized in a three-set Venn diagram to illustrate the unique and overlapping age-related variance captured by MEG, fMRI, and structural MRI.

###### General variance partitioning

We quantified the shared and unique variance among modalities using a three-step variance partitioning framework.

(1) Pairwise shared variance.

Each modality’s fingerprints were reduced to 50 principal components to ensure matched dimensionality and reduce overfitting. Pairwise linear regressions were performed between source and target fingerprints, and the coefficient of determination (𝑅^2^) quantified the variance in the target explained by the source.

(2) Pairwise CKA.

Linear CKA was computed between modalities to assess the correspondence of inter-subject similarity.

(3) Directional General Variance partitioning.

Using the same linear model setting as in (1), one of the three modalities (fMRI, MEG, or structural MRI) was set as the anchor, and its explained variance was decomposed into unique and shared components:

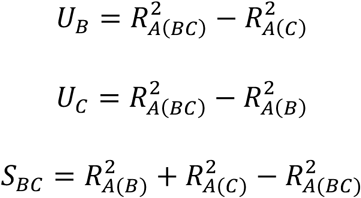

Partition results were visualized as three pie charts, each anchored on one modality, showing the relative proportions of unique and shared variance.

###### Cross-modal subject prediction

We are also interested in using a subject’s fingerprint extracted from one modality to predict the same subject’s fingerprint in the other modality. We extracted 𝑢, 𝑣 representing the fingerprints from two modalities. Within-modality subject prediction was relatively straightforward, as the source fingerprints and the target fingerprints share the same feature space. By contrast, cross-modality prediction is more complicated, as they are based on two different subject variability space. To map the source fingerprints 𝑢 to target fingerprints 𝑣 here, we need a projection between source fingerprints and target fingerprints. We first split the subjects into train set and test set (80:20). In train set, we trained a cross-modality projector 𝑓_Θ_: ℝ^𝑑𝑢^ → ℝ^𝑑𝑣^ to map source-modality fingerprints to target-modality fingerprints. Projector model parameters were optimized by minimizing the reconstruction loss:

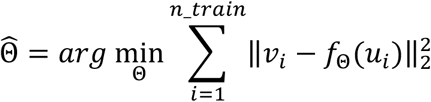

We tried a linear projector model and a non-linear model. For the linear model, 𝑓_Θ_(𝑢) = 𝑀𝑢, with 𝑀 ∈ ℝ^𝑑𝑢×𝑑𝑣^, the least-squares solution of 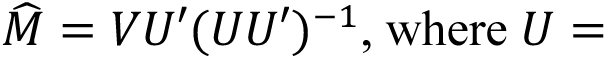 [𝑢_1_,…,𝑢_𝑛_𝑡𝑟𝑎𝑖𝑛_] and 𝑉 = [𝑣_1_,…, 𝑣_𝑛_𝑡𝑟𝑎𝑖𝑛_].

For the non-linear model, we adopted a single-layer multilayer perceptron (MLP) with ReLU activation function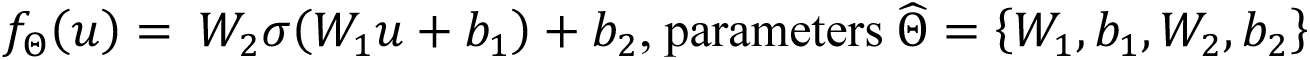 were optimized using gradient descent.

For each subject in the test set, projected target features from source were computed as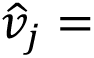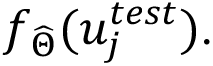 We then calculated similarity matrix 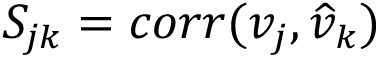 between projected and true target features; to estimate the subject identification accuracy 𝐴𝑐𝑐 between projected source fingerprints 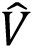 and target fingerprints 𝑉, we carried out the same approach as for the within-modality subject prediction described above. Additionally, we calculated the “top 5 accuracy”, defined as the proportion of test subjects for whom the true target fingerprint falls within the five candidates closest to the predicted fingerprint, to account for scenarios where a subject’s prediction is close but not exact, recognizing that some subjects may be closely clustered within the fingerprint space. To ensure robustness, the subjects were into train and test sets (80:20) five times randomly, repeating the process for each split. After all repetitions, we calculated the mean of the accuracy.

## 3. Results

### 3.1 Group-level correspondence of resting-state connectivity patterns

Group-level similarities in resting-state connectivity patterns have been reported in previous studies (Brookes et al., 2011; Hall et al., 2014). Consistent with these findings, as shown in Fig. 1, we observed a relatively strong correlation between the group-averaged static fMRI connectivity matrix and the static MEG connectivity matrices across frequency bands. This indicates that the large-scale network organization derived from fMRI and MEG shares a broadly similar topology at the group level. These results confirm that, although fMRI and MEG measure distinct aspects of brain activity, they converge on a similar spatial organization at the group level.

**Fig. 1.**
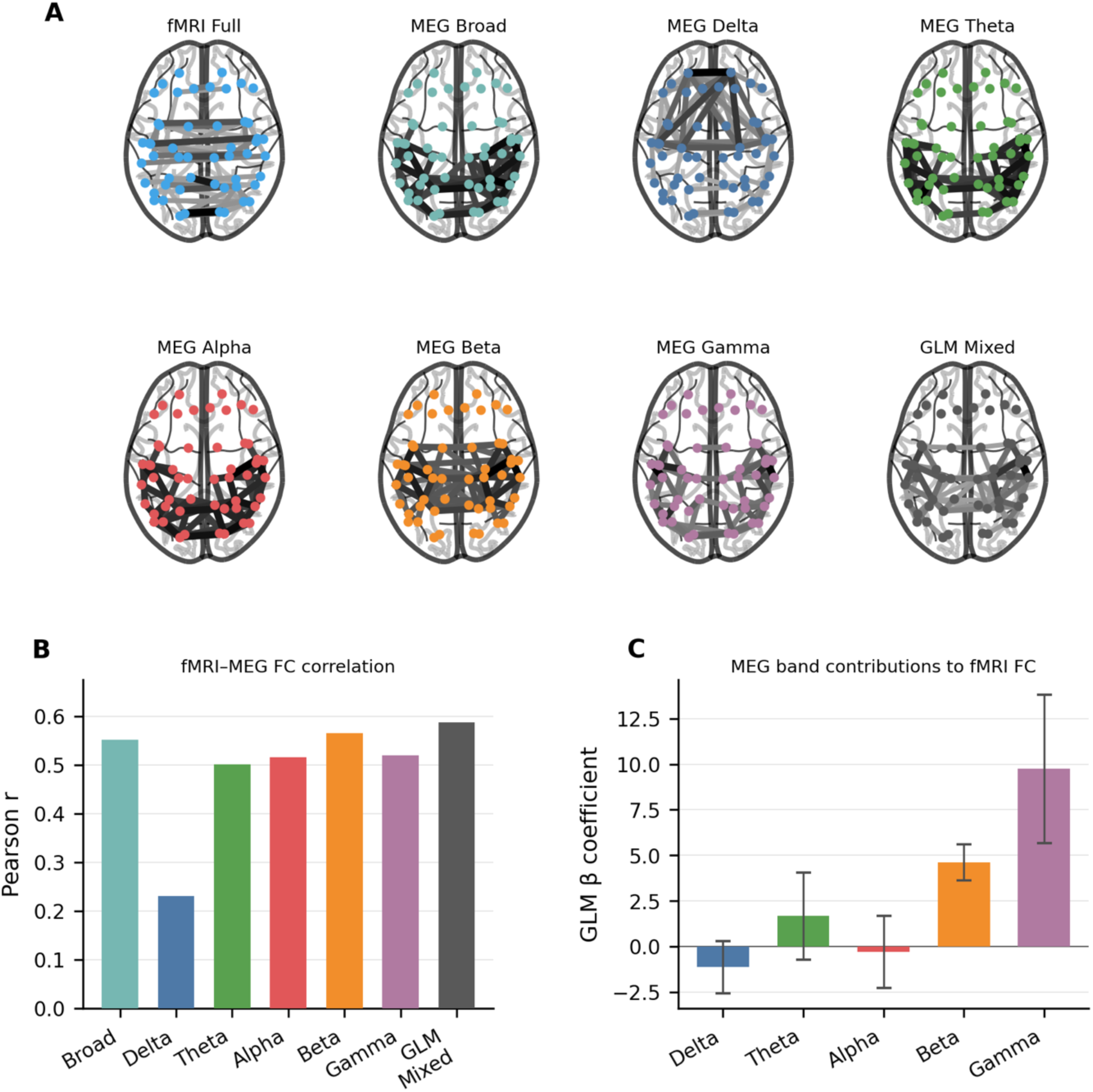
Group-level correspondence between time-averaged (static) MEG and fMRI FC. (A) **Glass brain visualisations** of group-averaged FC matrices for fMRI (full correlation) and MEG AEC across six frequency bands (broadband, delta, theta, alpha, beta, gamma), showing the top 5% strongest connections in the Glasser 52 parcellation. (B) **Correlation between MEG and fMRI group-level FC matrices.** Bar chart shows Pearson correlation coefficients between each MEG frequency band’s group-averaged FC and fMRI group-averaged FC, computed on the upper-triangle vectorised 52×52 matrices. The “GLM-mixed” bar represents the correlation achieved by combining all individual MEG frequency bands into a single general linear model (GLM) to predict fMRI FC. (C) **Contributions of individual frequency bands to the GLM-mixed prediction.** The GLM coefficients illustrate the relative weight of each MEG frequency band in driving the combined GLM-mixed result shown in panel B, with 95% confidence intervals from OLS standard errors.

### 3.2 fMRI and MEG yield robust, behaviourally relevant fingerprints

With both resting-state fMRI and MEG available for each subject in the Cam-CAN dataset, we estimated static fMRI, static MEG and dynamic MEG FCs for the same subjects using several widely adopted modelling approaches. These diverse methods were selected to comprehensively assess and compare their capacity to capture individual-level functional variability across modalities. We then extracted individual-level neural fingerprints from these estimated FCs to represent each subject across modalities and methods in a standardized feature space. For each feature, we evaluated the robustness of neural fingerprints by testing the discriminability and consistency across sessions.

Fig 2A illustrates the cross-session subject label prediction accuracy (i.e., discriminability) between sessions for all fingerprint features. Both static fMRI fingerprints, derived from time-averaged FC over the full scan, and dynamic MEG fingerprints, estimated from time-varying connectivity using time-delayed embedding (TDE) covariances or TDE-HMM state covariances, exhibited high differentiation accuracy, whereas static MEG fingerprints showed comparatively lower accuracy, indicating the presence of substantial within-subject variability across sessions. This within-subject, between-session variability may reflect either measurement-related factors (for example, head-position differences, scanner drift, and physiological artefacts) or genuine session-level fluctuation. The present design does not separate these sources, and the variability is therefore treated as nuisance variance in the subsequent fingerprint analyses (see Section 2.5.2.1).

**Fig. 2.**
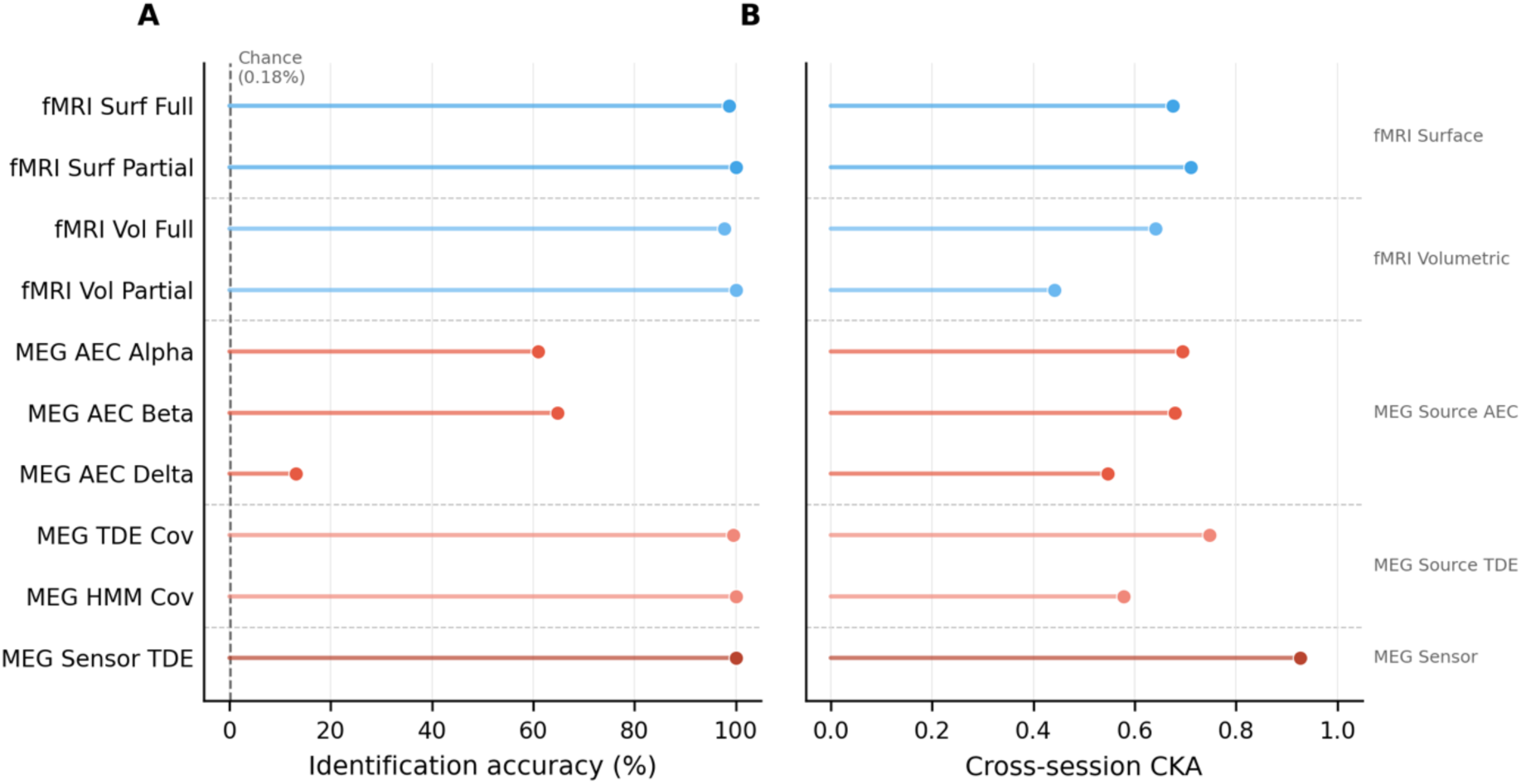
Cross-session robustness for different sets of fMRI and MEG fingerprints. (A) **Cross-session subject identification accuracy.** Each lollipop represents the percentage of correct subject identifications when matching individual fingerprints across sessions, demonstrating that fMRI-and MEG-derived features are generally reproducible across sessions. The vertical dashed line indicates chance level (0.18%, 1/543 subjects). (B) **Cross-session similarity measured by CKA.** High CKA values indicate preservation of the inter-subject similarity between different sessions of the data. Together, these results confirm that both fMRI and MEG fingerprints capture stable, individually distinctive neural patterns across sessions.

Beyond individual discriminability, we assessed the second-order test of whether the overall inter-subject similarity was preserved across sessions. To this end, we employed linear CKA, which compares representational similarity matrices while remaining invariant to coherent rotation and rescaling of the feature space. A fingerprint that captures genuine inter-subject similarity should therefore yield high CKA across sessions, even when absolute feature values drift. The resulting between-session CKA values are shown in Fig 2B. Most fingerprint features exhibit relatively high between-session consistency, indicating a stable inter-subject similarity across sessions. Sensor-space MEG features achieved the highest CKA values, followed by dynamic source-space MEG and static fMRI features, whereas static source-space MEG features, particularly at the gamma band, showed the lowest consistency (see Fig. S1). These rankings of discriminability and stability informed the feature selection for the cross-modal variance partitioning analysis presented in sections 3.4 and 3.5.

### 3.3 Neural fingerprints show meaningful variability regarding age and cognition

The preceding results establish that neural fingerprints are highly reproducible and unique to the individual. However, cross-session stability does not guarantee that these patterns are biologically informative. To determine whether this stable variability reflects meaningful differences, we next evaluated the predictive validity of these fingerprints. Specifically, we tested their ability to decode two key markers of individual diversity in the brain: chronological age and cognitive performance.

Fig 3A summarizes the age-prediction performance, reporting the coefficient of determination (𝑅^2^) between predicted and actual ages across test folds. An ElasticNet model was implemented within a 10-fold cross-validation scheme to obtain age predictions. For cognitive score prediction, 12 cognitive tasks were selected (Table 2) to maximize subject overlap with imaging data. Because age is a dominant source of variance that correlates with both neural fingerprints and cognitive performance, we first regressed out the effect of age from the cognitive scores to ensure that any subsequent brain-cognition associations reflect individual differences in cognition per se rather than shared age trends. PCA was then applied to these age-deconfounded cognitive scores to summarize shared and distinct sources of individual cognitive variability (Fig 3B). The first four components, each explaining >5% of the total variance, were retained for subsequent analysis. Principal Component 1 (PC1) was associated with fluid intelligence and executive function, whereas PC2 loaded broadly across memory, reasoning, emotion processing and reaction tasks, representing a general cognitive ability factor. PC3 was primarily associated with language processing, and PC4 captured variance related to language and memory.

**Fig. 3.**
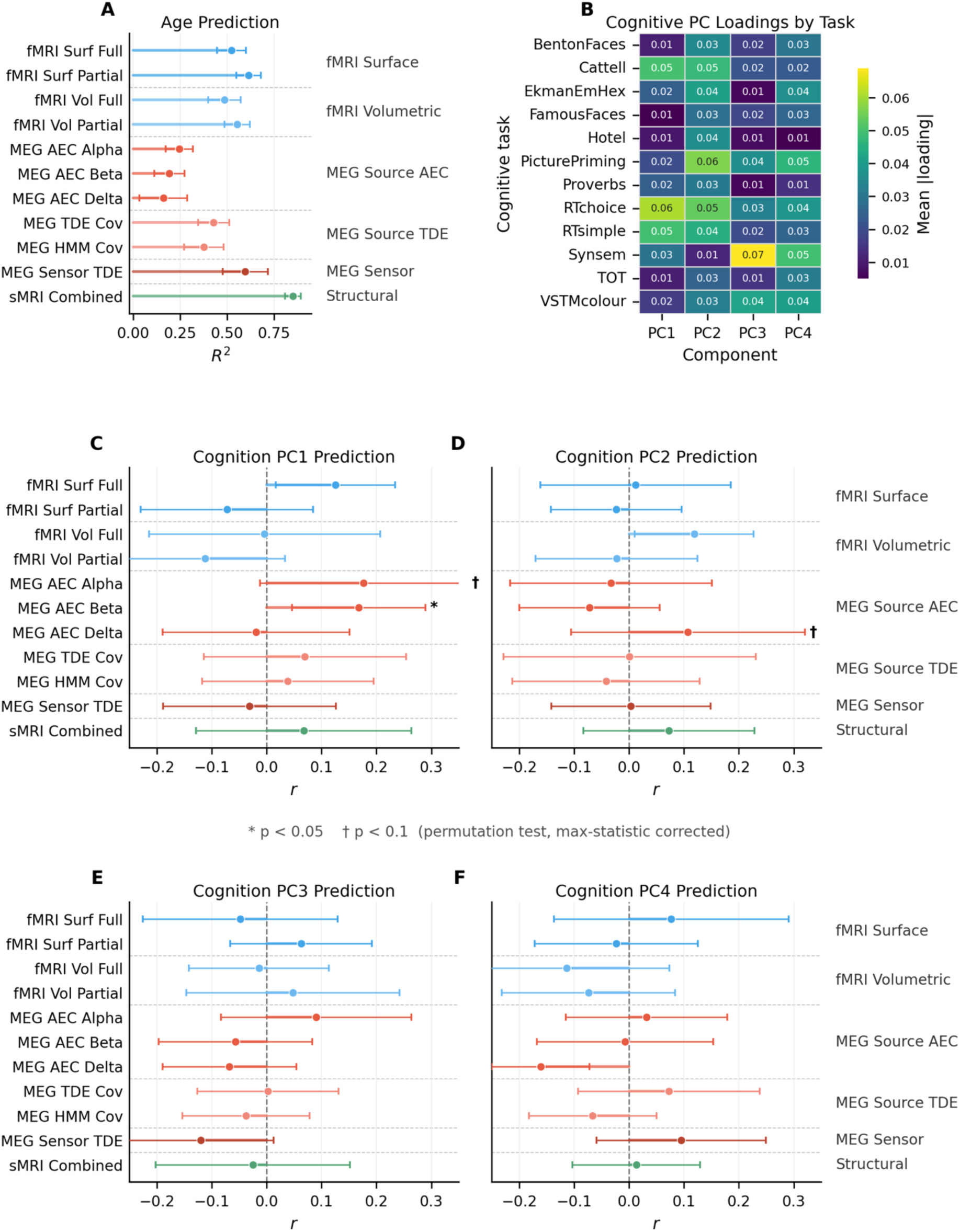
Age prediction and cognitive component prediction across modalities. (A) **Age prediction performance.** Horizontal lollipop plot showing cross-validated R² scores from 10-fold ElasticNet for each of the 11 main features, with error bars indicating standard deviation across folds. (B) **Cognitive component loadings.** Heatmap showing the mean absolute loadings of each cognitive task across the four principal components (PCs) (>5% variance explained) obtained from a PCA carried out on the age-deconfounded cognitive scores. (C–F) **Cross-validated cognitive prediction performance for PC1–PC4.** Pearson’s correlation (r) between predicted and observed component scores using 10-fold cross-validated ElasticNet. Statistical significance was determined by permutation testing, with maximum-statistic multiple comparison correction applied across all 26 features for each PC (* p<0.05, † p<0.1). PC1 showed the most robust prediction, driven mainly by static MEG fingerprints in the alpha and beta bands, while PC2 reached marginal significance (delta band), and PCs 3–4 were not predicted above chance.

We next evaluated the ability of the fingerprint features to predict each **cognitive** component. Using the same 10-fold cross-validated ElasticNet framework as in the age analysis, we quantified prediction performance as the Pearson’s r between predicted and observed component scores in the test sets. Statistical significance was determined using a 1000-iteration permutation test with maximum-statistic correction across all 26 features over each of the four components.

As shown in Fig 3C-F, PC1 was the most reliably predicted cognitive dimension, with significant correlations achieved by static MEG fingerprints in the alpha band (r=0.184, p=0.074, corrected) and beta band (r=0.191, p=0.035, corrected). Predictions for PC2 reached marginal significance, primarily driven by static MEG features in the delta band (r=0.159, p=0.087, corrected), while PC3 and PC4 were not predicted above chance by any modality.

Collectively, the age and cognitive prediction analyses demonstrate that these neural fingerprints capture biologically grounded individual variability. Age was robustly predicted across modalities, confirming the general sensitivity of the fingerprints to demographically driven variation. The cognitive results revealed more specific functional mappings. PC1 (fluid intelligence and executive function) was primarily driven by alpha (8–12 Hz) and beta (13–30 Hz) oscillations, consistent with their established roles in selective attention, information processing speed, and active cognitive regulation (Barone & Rossiter, 2021; Klimesch, 2012; Van Ede et al., 2010). In contrast, PC2 (memory and reasoning) was predominantly linked to delta oscillations (0.5–4 Hz), aligning with their known involvement in executive control and memory consolidation (Harmony, 2013). These frequency-specific associations are consistent with the established MEG literature, supporting the validity of the fingerprints as representations of functionally relevant individual differences.

It is also noteworthy that sensor-space MEG features outperformed source-space MEG features in both between-session robustness and prediction of age and cognitive components. This advantage likely arises because sensor-space representations retain individual-specific, non-neural factors confounds such as head size and positioning, which contribute additional variability. These factors are retained in sensor-space representations, are highly stable within subject across sessions, and partially covary with age; they therefore inflate between-session robustness and prediction of age and cognitive components without reflecting neurobiologically meaningful individual variability. In source space, leakage correction via symmetric orthogonalization is applied to increase confidence that the estimated connectivity reflects direct neuronal coupling rather than spatial artifact spread, but this suppression of shared variance can come at the expense of overall information content. Supporting this interpretation, supplementary analyses using source-space features without leakage correction (see Supplementary Information) revealed performance trends more similar to the sensor-space results, suggesting that the uncorrected features retain more total subject variability, albeit with reduced specificity to direct neuronal connectivity.

### 3.4 Structural factors mediate the age-related convergence of fMRI and MEG

The results so far demonstrate that both fMRI and MEG fingerprints are robust, individually unique, and sensitive to age. However, this parallel sensitivity to age does not by itself reveal whether the two modalities track the same underlying source of age-related variation. In particular, brain structure changes substantially with age and simultaneously constrains both hemodynamic and electrophysiological signals, raising the possibility that the apparent convergence between fMRI and MEG is driven by shared anatomical factors rather than reflecting direct functional coupling. To disentangle these contributions, we performed a variance partitioning analysis combining fMRI, MEG, and structural MRI (sMRI) features to isolate the unique and shared contributions of each modality to age prediction.

First, to enable cross-modal comparison, we selected a feature set for each modality that demonstrated strong cross-session robustness and predictive accuracy. For fMRI, we chose the surface-space *d50* partial correlation features, which consistently outperformed other metrics. For MEG, we selected the TDE covariance features, as they captured greater inter-individual variability than other static source-space MEG features, particularly in terms of between-session robustness and age prediction accuracy. Although these MEG features did not significantly predict cognition, likely because features capturing broad individual variability inherently contain more noise relative to subtle (and noisy) cognitive metrics, they offered the most robust electrophysiological fingerprint overall. For sMRI, we used a comprehensive feature set containing all the structural IDPs.

Second, we focused on age-related variance using variance partitioning, given that age represents a dominant source of individual-level variability in the Cam-CAN dataset. The results of this analysis are summarized in Table 4, which reports the predicted *R²* values for models including single modalities, pairwise combinations, and the full unified multimodal model. The seven variance components derived from these models were used to construct the Venn diagram in Fig 4, illustrating the unique and shared contributions of each modality to age-related variance.

**Fig. 4.**
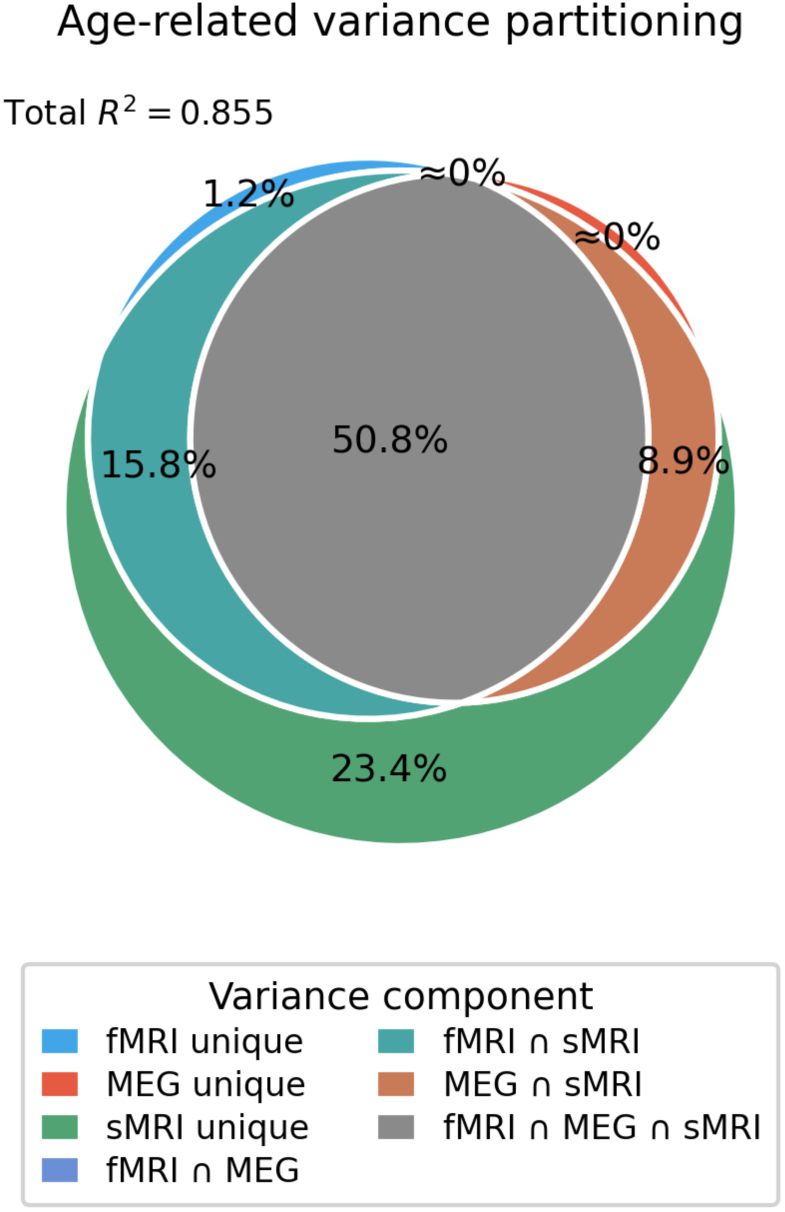
Age-related variance partitioning across modalities. Venn diagram illustrating the unique and shared age-related variance components among fMRI, MEG, and structural MRI fingerprints, based on inclusion-exclusion decomposition of cross-validated R² values. Structural features account for the largest portion of age-related variance (23.4% unique), with the majority of variance shared across all three modalities (50.8%). The fMRI∩MEG overlap and MEG unique are negligible (≈0%).

**Table 4:** Age-related variance partitioning results:

A: Age prediction results:
| Model | R<sup>2</sup> (age prediction) |
| --- | --- |
| A (fMRI) | 0.579 |
| B (MEG) | 0.483 |
| C (sMRI) | 0.845 |
| AB (fMRI + MEG) | 0.637 |
| AC (fMRI + sMRI) | 0.855 |
| BC (MEG + sMRI) | 0.818 |
| ABC (fMRI + MEG + sMRI) | 0.837 |

B: Variance partitioning interpreted results:
| Quantity | Interpreted R<sup>2</sup> |
| --- | --- |
| Triple overlap (fMRI ∩ MEG ∩ sMRI) | 0.434 |
| Overlap AB (fMRI ∩ MEG) | -0.009 |
| Overlap AC (fMRI ∩ sMRI) | 0.135 |
| Overlap BC (MEG ∩ sMRI) | 0.076 |
| Unique A (fMRI) | 0.010 |
| Unique B (MEG) | -0.027 |
| Unique C (sMRI) | 0.200 |

In Table 4, the variance partitioning results reveal small negative values in the fMRI-unique and MEG-fMRI shared components. This statistical artifact occurs when adding a redundant predictor (e.g., fMRI) to a dominant one (structure) increases model complexity without improving prediction, leading to a slight drop in generalization performance. This confirms that fMRI fingerprints provide virtually no unique age-related information beyond what is already captured by brain structure. Consequently, these negative values were treated as zero for the construction of the Venn diagram.

As shown in Fig 4, the variance shared by all three modalities was high. In contrast, the shared variance between fMRI and MEG was negligible, indicating that after controlling for structure, there is virtually no direct age-related overlap between the two functional modalities. In other words, functional age effects are largely explained by underlying structural changes rather than reflecting independent functional couplings.

### 3.5 Cross-modality similarities were found at individual level

While structural factors explain the shared age-related variance, we hypothesized that fMRI and MEG might still capture distinct sources of individual variability in other contexts. To quantify this direct correspondence, we attempted to predict subject identities across modalities (e.g., using an MEG fingerprint to identify the same subject in the fMRI dataset).

In contrast to the high within-modality reliability (Fig. 2), cross-modality identification was markedly lower. This limited direct correspondence confirms that, once structural constraints are accounted for, hemodynamic and electrophysiological fingerprints reflect largely non-redundant (i.e., unique) individual profiles.

To investigate this further, we examined the direct correspondence between fingerprint features across modalities using two complementary approaches. First, we performed shared variance subspace estimation to assess predictability, essentially asking: ‘can the MEG fingerprint of a given subject be predicted from the fMRI fingerprint of that same subject?’ This reveals the extent to which subject variability in one imaging domain can be directly inferred from another. Each cell in the heatmap in Fig. 5A indicates the proportion of variance in the target modality’s features that can be explained by the source modality.

**Fig. 5.**
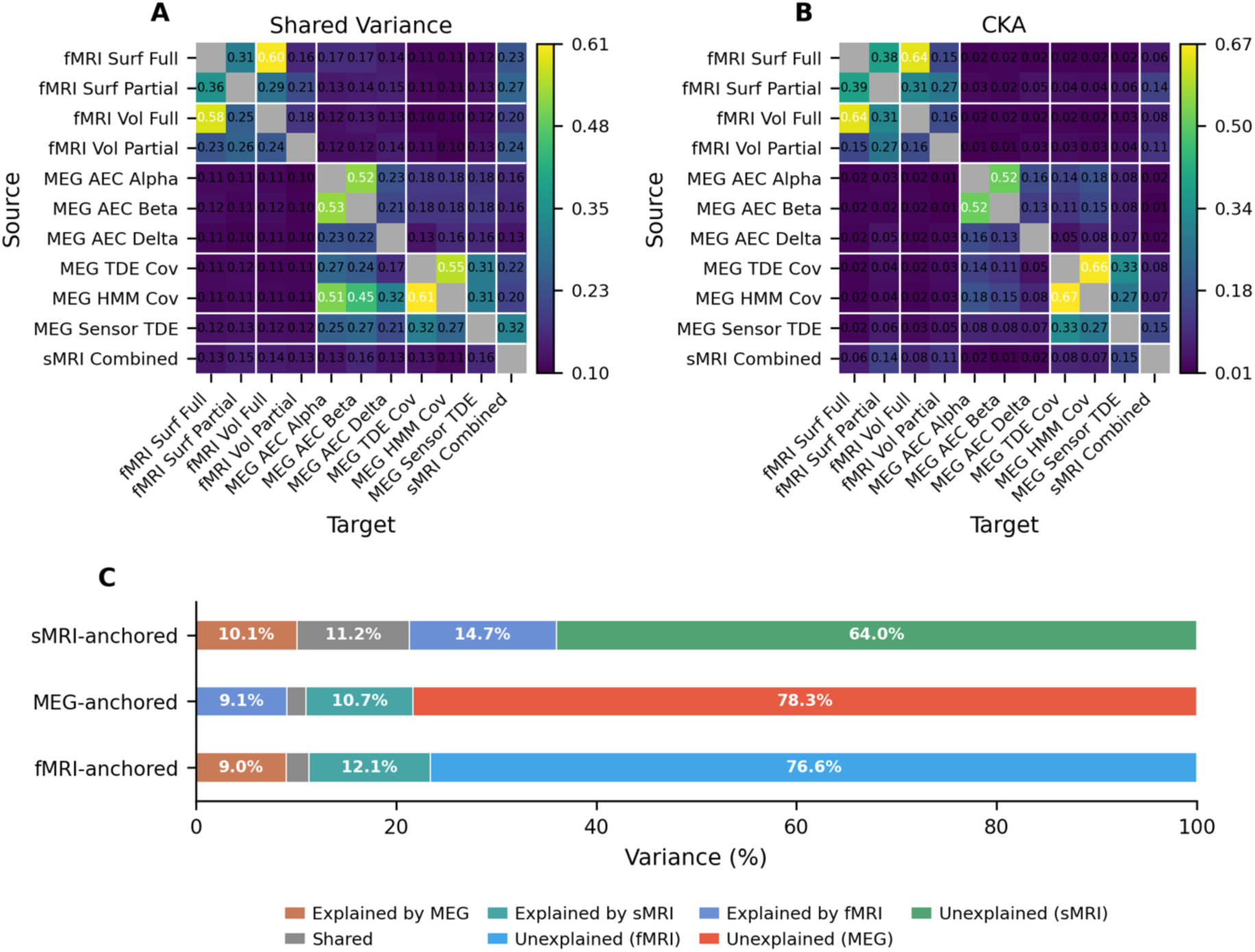
Individual-level variability is distinct across modalities. (A) **Cross-modal shared variance.** Heatmap (11×11) showing the proportion of variance in the source modality explained by the target modality via linear regression. Note the distinct modality-specific clustering, which contrasts with the high cross-modal correlation observed in group-averaged data (Fig. 1). (B) **Cross-modal similarities of inter-subject similarity (linear CKA).** Heatmap showing the similarities of inter-subject similarity between modalities, reflecting shared individual variability patterns beyond direct linear mapping. (C) **Directional general variance decomposition.** Each horizontal bar shows the decomposition of the variance for a different “anchor” modality into components explained by each of the other modalities and their overlap, alongside the unexplained (modality-specific) variance. Most variance is unexplained/modality-specific, while overlap between MEG-and fMRI-explained portions of structural variance indicates shared structural-related variability consistent with age-related variance partitioning results.

Second, we calculated linear CKA values to assess cross-modal correspondence of inter-subject similarity. This metric addresses a complementary question: ‘do subjects who appear similar in their fMRI scans also appear similar in their MEG scans?’, regardless of whether the specific features are directly predictable. The heatmap in Fig 5B presents the linear CKA values, which quantify the similarities of inter-subject similarity across fingerprint features.

Both heatmaps in Fig. 5 revealed clear clustering by modality. Within each modality, fingerprints exhibited high cross-feature similarity, indicating that features derived from the same modality capture largely consistent subject-specific representations. In contrast, cross-modal correspondence between MEG and fMRI was weak in both the shared subspace estimation and CKA analyses. This suggests that while a weak common signal exists, most of the individual variability is modality-specific, in contrast with the high cross-modal correlation observed in group-averaged data (see Fig. 1B).

Notably, moderate cross-modal associations were observed between structural and functional fingerprints, particularly for fMRI surface-space partial correlation features and TDE-MEG features. These relationships indicate that part of the subject variability in functional connectivity reflects structural constraints shared across modalities.

An asymmetric relationship was also evident in the shared variance heatmap (Fig. 5A): structural features could be partially inferred from functional features, whereas functional variability was poorly predicted by structural features. This imbalance suggests that, while structural organization imposes broad constraints that shape functional connectivity patterns, functional fingerprints capture additional subject variability that is not fully determined by anatomy.

To further characterize cross-modal relationships, we performed variance partitioning of the fingerprint features, summarized in Fig. 5C. Each chart illustrates how the variance of features from one anchored modality can be decomposed into components uniquely explained by that modality and those shared with the other two. Across modalities, a substantial proportion of variance was modality-specific, indicating that each fingerprint set captures considerable unique variability that is not shared with other modalities. This pattern is consistent with the clustering observed in the heatmaps (Fig. 5A–B), where within-modality similarity was markedly higher than cross-modality correspondence.

When partitioning the variance of structural features, we observed considerable overlap between the portions explained by MEG and by fMRI, suggesting that both functional modalities encode common structural-related variability. This finding aligns with the age-related variance partitioning results, reinforcing that MEG and fMRI share overlapping sensitivity to structural factors, while their direct functional overlap remains limited.

### 3.6 Cross-modality fingerprints prediction shows limited accuracy

Having observed at least a moderate degree of shared variance between modalities, we next assessed the extent to which a subject could be predicted (i.e., identified) across modalities. We used both a linear regression model and a nonlinear model implemented as a single-layer multilayer perceptron (MLP).

For each source–target feature pair, the model was trained to project fingerprint features from the source modality into the feature space of the target modality. Prediction accuracy was calculated in the test set as the proportion of subjects whose predicted target features were most similar to their true target features within the top 5 matches, thereby allowing tolerance for subjects with closely related fingerprint profiles.

The resulting cross-modal prediction accuracies are summarized as heatmaps in Fig. 6. The heatmaps revealed patterns broadly consistent with the shared-variance results. Within each modality, prediction accuracy was substantially higher than in cross-modality cases, forming clear modality-specific clusters. In contrast, cross-modal prediction accuracies were markedly lower, though still above the baseline, indicating a modest but nontrivial degree of shared individual variability across modalities.

**Fig. 6.**
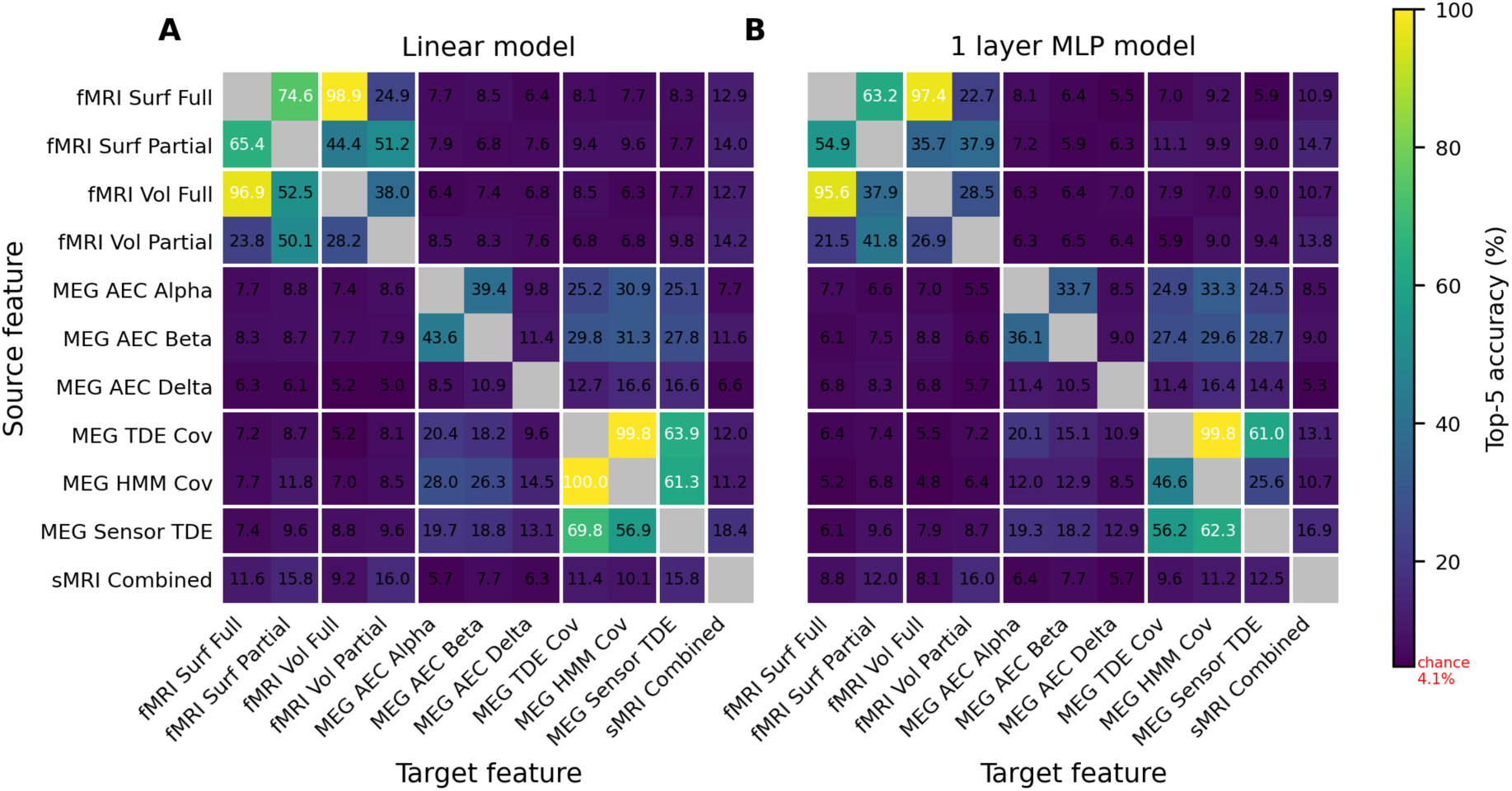
Cross-modal subject prediction performance. Heatmaps showing top-5 subject identification accuracy across all 11 main feature combinations using (A) a **linear model** and (B) a **single-layer MLP**. Each cell represents the percentage of correctly identified subjects in the test set (chance level = 4.10%, 5/122). Both models reveal above-chance cross-modal correspondence for selected feature pairs, with within-modality predictions substantially outperforming cross-modal predictions.

For fMRI, features based on full correlations showed strong mutual predictability across different parcellations and between volumetric and surface spaces, as did features based on partial correlations, suggesting that this variability is robust to spatial configuration.

However, prediction accuracy between full-and partial-correlation features was relatively low, indicating that they emphasize different aspects of subject variability.

For MEG, the amplitude envelope correlation (AEC) features exhibited reduced cross-band predictability, with lower accuracy between different frequency bands than between broadband and individual-band predictions. This pattern indicates that each frequency band captures distinct aspects of subject variability. Both static and dynamic features showed high mutual predictability, suggesting that these representations capture a consistent core of individual variability across temporal scales.

When comparing model performance, the linear model achieved slightly higher prediction accuracy than the single-layer MLP across most feature pairs. Given the current sample size, this result is consistent with a predominantly linear relationship between fingerprint features across modalities, though it does not rule out the possibility that nonlinear models could improve cross-modal correspondence given substantially more training data.

Consistent with the earlier variance partitioning results, age-related variance was well aligned across modalities, likely reflecting linear covariation driven by shared structural factors. In contrast, general variance partitioning revealed that only a small proportion of total variance was shared across modalities. Therefore, the low cross-modal prediction accuracy is better explained by the inherently limited shared subject variability between modalities than by shortcomings in model design or complexity.

## 4. Discussion

### 4.1 From group-level correspondence to individual-level complexity across modality

This study investigated the relationship between fMRI and MEG FC at the individual level. Previous group-level analyses have consistently reported strong correspondence of FC patterns across modalities, reflecting shared large-scale network organization. However, our results reveal that such cross-modal similarity becomes far more complex when examined at the individual-subject level.

While our results confirm that the underlying structure supports the group-level similarity of these functional modalities, this shared anatomy does not translate into a direct correspondence of functional fingerprints. This distinction highlights an important conceptual shift: group-level similarity does not imply similarity in how subjects vary. Variability in neural fingerprints arises from a combination of biological and methodological factors that vary differently across modalities and feature types, including anatomical constraints, hemodynamic response, electrophysiological oscillations, and subject-specific confounds. As a result, cross-modal relationships are more heterogeneous and less linearly aligned between fMRI and MEG than what is typically observed at the group level.

These findings reveal the complexity of analysing multimodal data at the individual level, where variability is inherently multidimensional rather than shared, and where each modality offers a complementary, rather than redundant, perspective on brain organization. This variability is also intertwined with subject-specific confounding factors, such as head motion, anatomical differences, or modality-specific noise sources, that are difficult to separate from ground-truth functional variability in neural fingerprints. Recognizing this complexity is essential for developing more precise individual-level models for MEG and fMRI, and for advancing multimodal fingerprinting approaches that account for both the shared and modality-specific information that each imaging technique contributes.

### 4.2 Structural dominance and its mediation of functional variability

Another important finding of this study is the dominant role of structural features in explaining age-related variance across modalities. Although both modalities individually predicted age with relatively high accuracy, their shared age-related variance was almost entirely explained by structural features. This finding suggests that the cross-modal convergence between fMRI and MEG arises not from direct correspondence between hemodynamic and electrophysiological coupling, but from mutual sensitivity to age-dependent structural variation (e.g., patterns of structural atrophy).

When examining the general variance partitioning, both functional modalities showed a moderate degree of overlap with structural features. This structural-functional overlap, while not as pronounced as in the age-related analysis, still indicates that structural organization remains an important source of subject variability across functional modalities. However, this does not imply that individual variability in functional fingerprints is dominated by structural factors. Indeed, we found that the unique variance within each functional modality still accounted for the largest proportion of total variability. The present decomposition cannot, however, distinguish whether this unique variance reflects genuine modality-specific subject variability or modality-specific noise.

### 4.3 Prediction of cognitive performance

Under the present analyses, only static MEG features successfully predicted cognitive performance above chance. A plausible contribution to this advantage is that MEG captures fast-scale oscillatory dynamics, particularly in the alpha beta, and delta bands, which are critical for executive function and working memory. These temporal features are not directly represented in sMRI or in the slower hemodynamic response of fMRI. The absence of significant cognitive prediction from fMRI and structural fingerprints here should not, however, be read as evidence that these modalities carry no cognitive information: earlier works have linked fMRI functional connectivity (Smith et al., 2015) and structural morphology (Llera et al., 2019) to behaviour or demographic measures in the Human Connectome Project (HCP). Two differences plausibly account for the discrepancy. First, the behavioural targets differ: the cognitive PCs used here summarise individual variability across all available Cam-CAN cognitive scores (N=375), whereas Smith et al. (2015) (N=461) and Llera et al. (2019) (N=448) related brain features to individual behavioural and demographic measures in the HCP. Second, the analytical criteria differ: the present study used 10-fold cross-validated ElasticNet regression onto each PC. Smith et al. (2015) instead performed a joint multivariate Canonical Correlation Analysis between resting-state functional connectome netmats and a large set of behavioural and demographic measures, identifying a single population-level mode of covariation rather than per-measure predictions. Llera et al. (2019) decomposed multi-modal structural features (including VBM, cortical thickness, and diffusion-derived indices) using Linked Independent Component Analysis and then tested simple linear correlations between each component’s subject loading and each behavioural or demographic measure separately. The present results therefore indicate that static MEG contributes cognitive information that is accessible under a strict, per-PC, held-out prediction criterion; whether fMRI and structural fingerprints would also predict cognition under this criterion using alternative behavioural targets or joint multivariate analyses remains an open question.

### 4.4 Capturing individual variability in MEG: TDE versus non-TDE features

We found that TDE-MEG features outperformed non-TDE AEC features in both cross-session fingerprint robustness and age prediction. This finding is consistent with previous reports showing that temporal autocorrelation features of MEG time series are most predictive of age (Stier et al., 2025), as TDE-MEG connectivity explicitly incorporates these temporal dependencies. TDE-MEG features capture richer temporal structure and, consequently, more subject-specific variability than non-TDE connectivity measures. In contrast, the cognition prediction results revealed better performance for non-TDE AEC features. This is likely due to the relatively small proportion of cognition-related variance present in resting-state data, combined with the higher signal-to-noise ratio (SNR) of frequency-band-specific AEC features for capturing cognitively relevant fluctuations.

Although TDE-MEG features encode broader aspects of subject variability, they may also include additional sources of variance that obscure the weaker cognition-related signal, thereby reducing prediction significance.

Together, these findings suggest that in MEG, TDE features provide a more comprehensive depiction of subject variability and are advantageous for capturing robust, temporally informed fingerprints. However, non-TDE AEC features retain value for detecting subtle behavioural effects under conditions where specific variance components, such as those linked to cognition, are comparatively small.

### 4.5 Static and dynamic MEG fingerprints: complementary but alike subject variability

We compared static TDE covariance features with dynamic features derived from TDE-HMM. Across all analyses, both static and dynamic TDE-MEG fingerprints showed comparable performance in cross-session robustness, age prediction, and cognition prediction. The absence of a clear advantage for dynamic features suggests that temporal dynamics do not contribute substantial additional subject variability beyond what is already captured by the time-delay embedding itself. Although the HMM framework models transient, state-specific fluctuations, these dynamics did not noticeably enhance subject identifiability or behavioural prediction.

### 4.6 Comparison of fMRI fingerprint features

We also compared fMRI fingerprints derived from surface-space (25-and 50-component) and volumetric-space parcellation (Glasser 52), using both full and partial correlations, to assess how spatial configuration and connectivity type influence subject variability.

In age prediction, partial correlation features outperformed full correlations, and surface-space representations achieved higher accuracy than volumetric ones. In fingerprint robustness, partial correlations again showed greater cross-session stability than full correlations, while surface-and volumetric-space features performed similarly. Regarding structure-function alignment, partial correlation features exhibited stronger correspondence with structural measures. In contrast, fingerprints remained highly consistent across parcellations regardless of whether full or partial correlations were used.

These distinctions reflect their differing statistical sensitivities: full correlations capture total associations, including indirect effects, whereas partial correlations isolate direct connections by accounting for shared variance with other regions. The stronger correspondence between partial-correlation fingerprints and structural features suggests that these direct functional connections are more tightly constrained by underlying anatomical architecture. Overall, the choice of correlation type exerts a greater influence on individual variability than the choice of spatial configuration, and surface-space parcellations offer a modest advantage for capturing subject variability.

### 4.7 The role of leakage correction in MEG source reconstruction

We also examined the effect of symmetric orthogonalization (Colclough et al., 2015), a leakage correction applied during MEG source reconstruction. As shown in the Supplementary Information, MEG features without leakage correction performed better in both age prediction and cross-session fingerprint robustness compared with leakage-corrected features. Similarly, sensor-space MEG features consistently outperformed source-space features in these analyses.

This pattern highlights an unintended consequence of leakage correction. While symmetric orthogonalization reduces artificial correlations caused by field spread and improves spatial specificity, it also removes subject-specific variance linked to stable but non-neural factors

such as head size, sensor positioning, and other acquisition-related artefacts. These factors contribute to inter-individual differences that enhance subject identifiability, even though they do not reflect neural activity of direct interest. Consequently, leakage-corrected source-space MEG features yield spatially cleaner but less individually distinctive fingerprints, whereas uncorrected or sensor-space features retain more subject-specific variability and achieve higher cross-session reproducibility.

Importantly, the geometry of the brain and head, which is itself a structural property, plays a fundamental role in shaping the MEG signal at the sensor level. Individual differences in cortical folding, skull thickness, and head shape directly influence the lead field and, therefore, the spatial distribution of measured magnetic fields. These structural factors inevitably introduce subject-specific variability into sensor-space data. Moreover, this structural influence cannot be eliminated even after source reconstruction, as forward models are imperfect approximations of the true volume conduction geometry. As a result, residual structural variability propagates into source-space estimates, meaning that neither sensor-space nor source-space MEG features are fully free from anatomical confounds. This interpretation is directly supported by supplementary directional general variance partitioning analyses in which the leakage-corrected MEG features were replaced by either uncorrected source-space or sensor-space MEG features (Supplementary Information). Compared with the leakage-corrected results, both alternative feature sets showed a higher proportion of structural variance explained by MEG and, symmetrically, a higher proportion of MEG variance explained by structural features, with the effect being most pronounced for sensor-space features. This graded pattern is consistent with the view that removing leakage correction, or moving to sensor space, progressively reintroduces structure-related subject variability into the MEG fingerprint. Notably, however, even after accounting for this structural contribution, a considerable portion of MEG variance remained modality-specific/unexplained across all feature variants, indicating that not all subject variability in MEG fingerprints can be attributed to structural factors alone.

This presents a critical trade-off for fingerprinting studies: maximizing subject identifiability via sensor-space or uncorrected features may come at the cost of biological specificity, as it retains stable non-neural confounds (e.g., head geometry). Future studies must therefore explicitly choose between optimizing for fingerprint strength versus biological fidelity, depending on their specific research goals.

This limitation also highlights a general challenge across all modalities: the unique variance of neural fingerprints remains difficult to disentangle. It is often unclear whether the extracted variability represents meaningful, functionally relevant individual differences or structured confounds inherent to data acquisition. Future work should aim to develop models that explicitly separate biologically meaningful variability from measurement-related and anatomical confounds, allowing neural fingerprints to more faithfully reflect genuine individual differences in brain organization.

### 4.8 Limitation

Several limitations of this study should be acknowledged. First, although the Cam-CAN dataset provides high-quality multimodal recordings, its sample size (N=543) is relatively modest for modelling complex, high-dimensional cross-modal relationships. This limits the feasibility of applying more advanced nonlinear or deep learning architectures for cross-modal alignment, which may require substantially larger datasets to avoid overfitting and ensure reliable generalization.

Second, the present analyses relied on conventional fingerprint modelling approaches for estimating session-specific covariance patterns. Emerging methods such as the HIVE (Hidden Markov Model with Integrated Variability Estimation)(Huang et al., 2026) could improve the precision of fingerprint estimation by jointly modelling both session-level and individual-level variability. Compared with standard HMM dual-estimation procedures, HIVE offers a more flexible representation of within-subject dynamics and may capture subtler interindividual differences.

Finally, all results were derived from resting-state data. Functional relationships between modalities, and the balance between shared and modality-specific variance, may differ under task conditions. Future studies incorporating task paradigms or longitudinal designs could help clarify how stable and context-dependent components of multimodal fingerprints jointly shape individual variability.

## Data and Code Availability

Raw data supporting the findings of this manuscript are available from the open-access Cambridge Centre for Ageing and Neuroscience (Cam-CAN) repository (Shafto et al., 2014; Taylor et al., 2017), can be requested at https://opendata.mrc-cbu.cam.ac.uk/projects/camcan/. The code used for MEG data analysis is accessible at https://github.com/OHBA-analysis/osl-dynamics. Structural and functional MRI preprocessing and analysis were conducted utilizing the UK Biobank pipeline, available at https://git.fmrib.ox.ac.uk/falmagro/UK_biobank_pipeline_v_1.

## Author Contributions

B.Z.M: Conceptualization, Methodology, Validation, Software, Data Curation, Formal Analysis, Writing - original draft, Visualization.

M.W.W: Conceptualization, Methodology, Data Curation, Supervision, Writing – reviewing & editing, Funding Acquisition.

S.M.S: Methodology, Supervision, Writing – reviewing & editing, Funding Acquisition.

## Ethics statement

Ethical approval for the original Cam-CAN study was granted by the Cambridgeshire 2 Research Ethics Committee (reference: 10/H0308/50). All participants provided written informed consent prior to participation in accordance with the Declaration of Helsinki. The current study constitutes a secondary analysis of these pre-existing, anonymized data.

## Acknowledgements

The authors thank Dr. Chetan Gohil for preprocessing the MEG data, and Dr. Christine Ahrends, Dr. Gaurav Bhalerao and Dr. Fidel Alfaro-Almagro for their assistance in setting up the UK Biobank pipeline for the Cam-CAN dataset. We are grateful to Professor Richard N. Henson at the University of Cambridge for his help in facilitating access to the Cam-CAN dataset and associated MRI gradient coefficient files. We also acknowledge the members of OxCIN for their constructive discussions during this project.

## Reference

Alfaro-Almagro, F., Jenkinson, M., Bangerter, N. K., Andersson, J. L. R., Griffanti, L., Douaud, G., Sotiropoulos, S. N., Jbabdi, S., Hernandez-Fernandez, M., Vallee, E., Vidaurre, D., Webster, M., McCarthy, P., Rorden, C., Daducci, A., Alexander, D. C., Zhang, H., Dragonu, I., Matthews, P. M.,…Smith, S. M. (2018). Image processing and Quality Control for the first 10,000 brain imaging datasets from UK Biobank. NeuroImage, 166, 400–424. 10.1016/j.neuroimage.2017.10.034

Baker, A. P., Brookes, M. J., Rezek, I. A., Smith, S. M., Behrens, T., Probert Smith, P. J., & Woolrich, M. (2014). Fast transient networks in spontaneous human brain activity. eLife, 3, e01867. 10.7554/eLife.01867

Barone, J., & Rossiter, H. E. (2021). Understanding the Role of Sensorimotor Beta Oscillations. Frontiers in Systems Neuroscience, 15, 655886. 10.3389/fnsys.2021.655886

Beckmann, C. F., & Smith, S. M. (2004). Probabilistic Independent Component Analysis for Functional Magnetic Resonance Imaging. IEEE Transactions on Medical Imaging, 23(2), 137–152. 10.1109/TMI.2003.822821

Bijsterbosch, J. D., Woolrich, M. W., Glasser, M. F., Robinson, E. C., Beckmann, C. F., Van Essen, D. C., Harrison, S. J., & Smith, S. M. (2018). The relationship between spatial configuration and functional connectivity of brain regions. eLife, 7, e32992. 10.7554/eLife.32992

Biswal, B., Zerrin Yetkin, F., Haughton, V. M., & Hyde, J. S. (1995). Functional connectivity in the motor cortex of resting human brain using echo-planar mri. Magnetic Resonance in Medicine, 34(4), 537–541. 10.1002/mrm.1910340409

Brookes, M. J., Woolrich, M., Luckhoo, H., Price, D., Hale, J. R., Stephenson, M. C., Barnes, G. R., Smith, S. M., & Morris, P. G. (2011). Investigating the electrophysiological basis of resting state networks using magnetoencephalography. Proceedings of the National Academy of Sciences, 108(40), 16783–16788. 10.1073/pnas.1112685108

Cho, S., Van Es, M., Woolrich, M., & Gohil, C. (2024). Comparison between EEG and MEG of static and dynamic resting-state networks. Human Brain Mapping, 45(13), e70018. 10.1002/hbm.70018

Colclough, G. L., Brookes, M. J., Smith, S. M., & Woolrich, M. W. (2015). A symmetric multivariate leakage correction for MEG connectomes. NeuroImage, 117, 439–448. 10.1016/j.neuroimage.2015.03.071

Colclough, G. L., Woolrich, M. W., Tewarie, P. K., Brookes, M. J., Quinn, A. J., & Smith, S. M. (2016). How reliable are MEG resting-state connectivity metrics? NeuroImage, 138, 284–293. 10.1016/j.neuroimage.2016.05.070

De Pasquale, F., Della Penna, S., Snyder, A. Z., Lewis, C., Mantini, D., Marzetti, L., Belardinelli, P., Ciancetta, L., Pizzella, V., Romani, G. L., & Corbetta, M. (2010). Temporal dynamics of spontaneous MEG activity in brain networks. Proceedings of the National Academy of Sciences, 107(13), 6040–6045. 10.1073/pnas.0913863107

Fox, M. D., & Raichle, M. E. (2007). Spontaneous fluctuations in brain activity observed with functional magnetic resonance imaging. Nature Reviews Neuroscience, 8(9), 700–711. 10.1038/nrn2201

Gohil, C., Huang, R., Roberts, E., Van Es, M. W., Quinn, A. J., Vidaurre, D., & Woolrich, M. W. (2024). Osl-dynamics, a toolbox for modeling fast dynamic brain activity. eLife, 12, RP91949. 10.7554/eLife.91949.3

Griffanti, L., Douaud, G., Bijsterbosch, J., Evangelisti, S., Alfaro-Almagro, F., Glasser, M. F., Duff, E. P., Fitzgibbon, S., Westphal, R., Carone, D., Beckmann, C. F., & Smith, S. M. (2017). Hand classification of fMRI ICA noise components. NeuroImage, 154, 188–205. 10.1016/j.neuroimage.2016.12.036

Griffanti, L., Salimi-Khorshidi, G., Beckmann, C. F., Auerbach, E. J., Douaud, G., Sexton, C. E., Zsoldos, E., Ebmeier, K. P., Filippini, N., Mackay, C. E., Moeller, S., Xu, J., Yacoub, E., Baselli, G., Ugurbil, K., Miller, K. L., & Smith, S. M. (2014). ICA-based artefact removal and accelerated fMRI acquisition for improved resting state network imaging. NeuroImage, 95, 232–247. 10.1016/j.neuroimage.2014.03.034

Hall, E. L., Robson, S. E., Morris, P. G., & Brookes, M. J. (2014). The relationship between MEG and fMRI. NeuroImage, 102, 80–91. 10.1016/j.neuroimage.2013.11.005

Harmony, T. (2013). The functional significance of delta oscillations in cognitive processing. Frontiers in Integrative Neuroscience, 7. 10.3389/fnint.2013.00083

Huang, R., Gohil, C., & Woolrich, M. (2026). Modelling variability in functional brain networks using embeddings. Imaging Neuroscience. 10.1162/IMAG.a.1188

Jenkinson, M., Bannister, P., Brady, M., & Smith, S. (2002). Improved Optimization for the Robust and Accurate Linear Registration and Motion Correction of Brain Images. NeuroImage, 17(2), 825–841. 10.1006/nimg.2002.1132

Klimesch, W. (2012). Alpha-band oscillations, attention, and controlled access to stored information. Trends in Cognitive Sciences, 16(12), 606–617. 10.1016/j.tics.2012.10.007

Kohl, O., Woolrich, M., Nobre, A. C., & Quinn, A. (2023). *Glasser52: A parcellation for MEG-Analysis* [Data set]. Zenodo. 10.5281/ZENODO.10401792

Llera, A., Wolfers, T., Mulders, P., & Beckmann, C. F. (2019). Inter-individual differences in human brain structure and morphology link to variation in demographics and behavior. eLife, 8, e44443. 10.7554/eLife.44443

Mantini, D., Perrucci, M. G., Del Gratta, C., Romani, G. L., & Corbetta, M. (2007). Electrophysiological signatures of resting state networks in the human brain. Proceedings of the National Academy of Sciences, 104(32), 13170–13175. 10.1073/pnas.0700668104

Quinn, A. J., van Es, M. W. J., Gohil, C., & Woolrich, M. W. (2022). OHBA Software Library in Python (OSL) (Version 0.1.1) [Computer software]. Zenodo. 10.5281/ZENODO.6875060

Ray-Mukherjee, J., Nimon, K., Mukherjee, S., Morris, D. W., Slotow, R., & Hamer, M. (2014). Using commonality analysis in multiple regressions: A tool to decompose regression effects in the face of multicollinearity. Methods in Ecology and Evolution, 5(4), 320–328. 10.1111/2041-210X.12166

Salimi-Khorshidi, G., Douaud, G., Beckmann, C. F., Glasser, M. F., Griffanti, L., & Smith, S. M. (2014). Automatic denoising of functional MRI data: Combining independent component analysis and hierarchical fusion of classifiers. NeuroImage, 90, 449–468. 10.1016/j.neuroimage.2013.11.046

Seibold, D. R., & McPHEE, R. D. (1979). COMMONALITY ANALYSIS: A METHOD FOR DECOMPOSING EXPLAINED VARIANCE IN MULTIPLE REGRESSION ANALYSES. Human Communication Research, 5(4), 355–365. 10.1111/j.1468-2958.1979.tb00649.x

Shafto, M. A., Dixon, M., Taylor, J. R., Rowe, J. B., Cusack, R., Calder, A. J., Marslen-Wilson, W. D., Duncan, J., Dalgleish, T., Henson, R. N., Brayne, C., & Matthews, F. E. (2014). The Cambridge Centre for Ageing and Neuroscience (Cam-CAN) study protocol: A cross-sectional, lifespan, multidisciplinary examination of healthy cognitive ageing. BMC Neurology, 14(1), 204. 10.1186/s12883-014-0204-1

Shulman, G. L., Fiez, J. A., Corbetta, M., Buckner, R. L., Miezin, F. M., Raichle, M. E., & Petersen, S. E. (1997). Common Blood Flow Changes across Visual Tasks: II. Decreases in Cerebral Cortex. Journal of Cognitive Neuroscience, 9(5), 648–663. 10.1162/jocn.1997.9.5.648

Smith, S. M., Nichols, T. E., Vidaurre, D., Winkler, A. M., Behrens, T. E. J., Glasser, M. F., Ugurbil, K., Barch, D. M., Van Essen, D. C., & Miller, K. L. (2015). A positive-negative mode of population covariation links brain connectivity, demographics and behavior. Nature Neuroscience, 18(11), 1565–1567. 10.1038/nn.4125

Stier, C., Balestrieri, E., Fehring, J., Focke, N. K., Wollbrink, A., Dannlowski, U., & Gross, J. (2025). Temporal autocorrelation is predictive of age—An extensive MEG time-series analysis. Proceedings of the National Academy of Sciences, 122(8), e2411098122. 10.1073/pnas.2411098122

Taylor, J. R., Williams, N., Cusack, R., Auer, T., Shafto, M. A., Dixon, M., Tyler, L. K., Cam-CAN, & Henson, R. N. (2017). The Cambridge Centre for Ageing and Neuroscience (Cam-CAN) data repository: Structural and functional MRI, MEG, and cognitive data from a cross-sectional adult lifespan sample. NeuroImage, 144, 262–269. 10.1016/j.neuroimage.2015.09.018

Van Ede, F., Jensen, O., & Maris, E. (2010). Tactile expectation modulates pre-stimulus β-band oscillations in human sensorimotor cortex. NeuroImage, 51(2), 867–876. 10.1016/j.neuroimage.2010.02.053

Van Es, M. W. J., Gohil, C., Quinn, A. J., & Woolrich, M. W. (2025). osl-ephys: A Python toolbox for the analysis of electrophysiology data. Frontiers in Neuroscience, 19, 1522675. 10.3389/fnins.2025.1522675

Van Veen, B. D., & Buckley, K. M. (1988). Beamforming: A versatile approach to spatial filtering. IEEE ASSP Magazine, 5(2), 4–24. 10.1109/53.665

Vidaurre, D., Hunt, L. T., Quinn, A. J., Hunt, B. A. E., Brookes, M. J., Nobre, A. C., & Woolrich, M. W. (2018). Spontaneous cortical activity transiently organises into frequency specific phase-coupling networks. Nature Communications, 9(1), 2987. 10.1038/s41467-018-05316-z

