## Supplementary Information (SI) for "fMRI and MEG Fingerprints Diverge at the Individual Level"

A. All cognitive scores

| Task | scores |
| --- | --- |
| BentonFaces | BentonFaces_SubScore1; BentonFaces_SubScore2; BentonFaces_TotalScore |
| Cattell | Cattell_SubScore1; Cattell_SubScore2; Cattell_SubScore3; Cattell_SubScore4; Cattell_TotalScore |
| EkmanEmHex | EkmanEmHex_Ang_Corr; EkmanEmHex_Ang_Acc; EkmanEmHex_Ang_Ang; EkmanEmHex_Dis_Corr; EkmanEmHex_Dis_Acc; EkmanEmHex_Dis_Dis; EkmanEmHex_Fea_Corr; EkmanEmHex_Fea_Acc; EkmanEmHex_Fea_Fea; EkmanEmHex_Hap_Corr; EkmanEmHex_Hap_Acc; EkmanEmHex_Hap_Hap; EkmanEmHex_Sad_Corr; EkmanEmHex_Sad_Acc; EkmanEmHex_Sad_Sad; EkmanEmHex_Sur_Corr; EkmanEmHex_Sur_Acc; EkmanEmHex_Sur_Sur |
| FamousFaces | FamousFaces_FacesTest_FAMnam; FamousFaces_FacesTest_FAMocc; FamousFaces_FacesTest_FAMfam; FamousFaces_FacesTest_FAMunfam; FamousFaces_FacesTest_UNnam; FamousFaces_FacesTest_UNocc; FamousFaces_FacesTest_UNfam; FamousFaces_FacesTest_UNunfam; FamousFaces_NamesTest_FAMnam; FamousFaces_NamesTest_FAMocc; FamousFaces_NamesTest_FAMfam; FamousFaces_NamesTest_FAMunfam; FamousFaces_NamesTest_UNnam; FamousFaces_NamesTest_UNocc; FamousFaces_NamesTest_UNfam; FamousFaces_NamesTest_UNunfam |
| Hotel | Hotel_Time |
| PicturePriming | PicturePriming_ncorrect; PicturePriming_nincorrect; PicturePriming_ndontknow; PicturePriming_nhesitation; PicturePriming_nnoresp; PicturePriming_ncheckresp; PicturePriming_nMic0Resp0; PicturePriming_nMic1Resp0; PicturePriming_nMic0Resp1; PicturePriming_nMic1Resp1; PicturePriming_nRTearly; PicturePriming_nRTusable; PicturePriming_nRTcorrect; PicturePriming_ACC_baseline_all; PicturePriming_ACC_priming_all; PicturePriming_reITRT_baseline_all; PicturePriming_reITRT_priming_target_all; PicturePriming_reITRT_priming_prime_all |
| Proverbs | Proverbs_Score |
| RTchoice | RTchoice_PctCorrect_all; RTchoice_RTmean_all; RTchoice_RTmedian_all; RTchoice_RTsd_all; RTchoice_RTcv_all; RTchoice_RTmean_73; RTchoice_RTmedian_73; RTchoice_RTsd_73; RTchoice_RTcv_73; RTchoice_RTmean_79; RTchoice_RTmedian_79; RTchoice_RTsd_79; RTchoice_RTcv_79; RTchoice_RTmean_85; RTchoice_RTmedian_85; RTchoice_RTsd_85; RTchoice_RTcv_85; RTchoice_RTmean_59; RTchoice_RTmedian_59; RTchoice_RTsd_59; RTchoice_RTcv_59 |
| RTsimple | RTsimple_PctCorrect; RTsimple_RTmean; RTsimple_RTmedian; RTsimple_RTsd; RTsimple_RTcv; RTsimple_invRTmean; RTsimple_invRTmedian; RTsimple_invRTsd; RTsimple_invRTcv |
| Synsem | Synsem_synsub_RT_mean; Synsem_syndom_RT_mean; Synsem_synamb_RT_mean; Synsem_synunamb_RT_mean; Synsem_synunacc_RT_mean; Synsem_semsub_RT_mean; Synsem_semdom_RT_mean; Synsem_semamb_RT_mean; Synsem_semunamb_RT_mean; Synsem_semunacc_RT_mean; Synsem_synsub_ERR_mean; Synsem_syndom_ERR_mean; Synsem_synamb_ERR_mean; Synsem_synunamb_ERR_mean; Synsem_synunacc_ERR_mean; Synsem_semsub_ERR_mean; Synsem_semdom_ERR_mean; Synsem_semamb_ERR_mean; Synsem_semunamb_ERR_mean; Synsem_semunacc_ERR_mean |
| TOT | TOT_sum_KC; TOT_sum_KI; TOT_sum_TO; TOT_sum_DK; TOT_sum_SE; TOT_sum_NU; TOT_ToT_ratio |
| VSTMcolour | VSTMcolour_Prcsn_ss1; VSTMcolour_K_ss1; VSTMcolour_Doubt_ss1; VSTMcolour_RT_ss1; VSTMcolour_MSE_ss1; VSTMcolour_Prcsn_ss2; VSTMcolour_K_ss2; VSTMcolour_Doubt_ss2; VSTMcolour_NonTarg_ss2; VSTMcolour_RT_ss2; VSTMcolour_MSE_ss2; VSTMcolour_Prcsn_ss3; VSTMcolour_K_ss3; VSTMcolour_Doubt_ss3; VSTMcolour_NonTarg_ss3; VSTMcolour_RT_ss3; VSTMcolour_MSE_ss3; VSTMcolour_Prcsn_ss4; VSTMcolour_K_ss4; VSTMcolour_Doubt_ss4; VSTMcolour_NonTarg_ss4; VSTMcolour_RT_ss4; VSTMcolour_MSE_ss4 |

B. full cognitive prediction results:

| PC | Feature | Observed Mean r (corrected p value) |
| --- | --- | --- |
| 1 | nc_aec_alpha | 0.215 (p=0.0320) |
| 1 | aec_beta | 0.191 (p=0.0350) |
| 1 | aec_alpha | 0.184 (p=0.0749) |
| 2 | aec_delta | 0.159 (p=0.0869) |
| 1 | nc_aec_beta | 0.170 (p=0.1249) |
| 1 | hmm_covs | 0.136 (p=0.1429) |
| 1 | aec_broad | 0.162 (p=0.1738) |
| 1 | nc_aec_broad | 0.142 (p=0.2637) |
| 2 | fmri_parcel_full | 0.139 (p=0.4296) |
| 2 | nc_parcel_tde_cov | 0.131 (p=0.5325) |
| 2 | nc_aec_delta | 0.105 (p=0.5754) |
| 1 | fmri_surface_d50_full | 0.131 (p=0.5754) |
| 1 | parcel_tde_cov | 0.118 (p=0.6124) |
| 4 | parcel_tde_cov | 0.108 (p=0.6284) |
| 2 | nc_aec_gamma | 0.101 (p=0.6733) |
| 1 | nc_parcel_tde_cov | 0.103 (p=0.7163) |
| 2 | nc_aec_beta | 0.088 (p=0.7233) |
| 1 | fmri_surface_d25_full | 0.127 (p=0.7293) |
| 1 | structural_concat | 0.088 (p=0.7572) |
| 4 | fmri_surface_d25_full | 0.118 (p=0.7732) |
| 2 | fmri_surface_d25_partial | 0.116 (p=0.8212) |
| 2 | aec_theta | 0.087 (p=0.8282) |
| 4 | nc_aec_alpha | 0.108 (p=0.8372) |
| 3 | aec_beta | 0.081 (p=0.8472) |
| 3 | fmri_surface_d25_full | 0.104 (p=0.8621) |
| 4 | aec_gamma | 0.094 (p=0.8751) |
| 4 | aec_theta | 0.095 (p=0.8751) |
| 1 | fmri_surface_d50_partial | 0.084 (p=0.8771) |
| 1 | aec_theta | 0.072 (p=0.8911) |
| 4 | sensor_tde_cov | 0.072 (p=0.9191) |
| 4 | nc_aec_beta | 0.087 (p=0.9381) |
| 4 | nc_parcel_tde_cov | 0.064 (p=0.9580) |
| 4 | fmri_surface_d50_partial | 0.070 (p=0.9770) |
| 2 | structural_FS | 0.057 (p=0.9840) |
| 4 | aec_broad | 0.061 (p=0.9840) |
| 3 | aec_delta | 0.053 (p=0.9870) |
| 3 | nc_aec_broad | 0.053 (p=0.9890) |
| 2 | fmri_surface_d50_full | 0.046 (p=0.9950) |
| 4 | structural_concat | 0.048 (p=0.9960) |
| 2 | hmm_covs | 0.048 (p=0.9960) |
| 2 | aec_gamma | 0.039 (p=0.9970) |
| 1 | sensor_tde_cov | 0.030 (p=0.9980) |
| 1 | aec_delta | 0.046 (p=0.9980) |
| 2 | sensor_tde_cov | 0.048 (p=0.9980) |
| 1 | nc_aec_gamma | 0.028 (p=0.9990) |
| 2 | fmri_surface_d50_partial | 0.034 (p=0.9990) |
| 1 | nc_aec_delta | 0.034 (p=0.9990) |
| 1 | aec_gamma | -0.009 (p=1.0000) |
| 4 | structural_FS | 0.002 (p=1.0000) |
| 3 | fmri_surface_d50_full | -0.082 (p=1.0000) |
| 2 | nc_aec_alpha | -0.078 (p=1.0000) |
| 1 | fmri_surface_d25_partial | 0.031 (p=1.0000) |
| 4 | nc_aec_broad | -0.013 (p=1.0000) |
| 4 | nc_aec_gamma | -0.025 (p=1.0000) |
| 3 | nc_parcel_tde_cov | -0.083 (p=1.0000) |
| 3 | fmri_surface_d25_partial | -0.029 (p=1.0000) |
| 2 | aec_alpha | -0.116 (p=1.0000) |
| 2 | sensor_cov | -0.038 (p=1.0000) |
| 3 | fmri_parcel_full | -0.037 (p=1.0000) |
| 3 | nc_aec_beta | -0.002 (p=1.0000) |
| 4 | aec_alpha | 0.008 (p=1.0000) |
| 3 | sensor_cov | -0.066 (p=1.0000) |
| 1 | structural_FS | -0.088 (p=1.0000) |
| 2 | parcel_tde_cov | -0.021 (p=1.0000) |
| 4 | fmri_parcel_full | -0.062 (p=1.0000) |
| 3 | aec_broad | 0.007 (p=1.0000) |
| 2 | fmri_parcel_partial | 0.016 (p=1.0000) |
| 3 | aec_alpha | -0.075 (p=1.0000) |
| 2 | structural | 0.018 (p=1.0000) |
| 3 | fmri_surface_d50_partial | -0.017 (p=1.0000) |
| 2 | aec_broad | 0.016 (p=1.0000) |
| 4 | hmm_covs | -0.025 (p=1.0000) |
| 4 | fmri_surface_d50_full | 0.010 (p=1.0000) |
| 4 | sensor_cov | 0.024 (p=1.0000) |
| 3 | nc_aec_delta | -0.026 (p=1.0000) |
| 3 | aec_theta | -0.049 (p=1.0000) |
| 2 | aec_beta | -0.013 (p=1.0000) |
| 2 | fmri_surface_d25_full | 0.020 (p=1.0000) |
| 2 | nc_aec_broad | -0.015 (p=1.0000) |
| 4 | nc_aec_theta | -0.132 (p=1.0000) |
| 4 | aec_delta | -0.042 (p=1.0000) |
| 3 | parcel_tde_cov | -0.012 (p=1.0000) |
| 2 | structural_concat | -0.037 (p=1.0000) |
| 3 | nc_aec_alpha | -0.004 (p=1.0000) |
| 3 | structural_concat | -0.053 (p=1.0000) |
| 3 | nc_aec_gamma | 0.008 (p=1.0000) |
| 4 | fmri_parcel_partial | -0.016 (p=1.0000) |
| 3 | aec_gamma | -0.066 (p=1.0000) |
| 3 | nc_aec_theta | 0.027 (p=1.0000) |
| 1 | structural | -0.071 (p=1.0000) |
| 1 | nc_aec_theta | -0.016 (p=1.0000) |
| 1 | fmri_parcel_full | -0.037 (p=1.0000) |
| 3 | fmri_parcel_partial | 0.035 (p=1.0000) |
| 1 | sensor_cov | -0.015 (p=1.0000) |
| 4 | structural | 0.024 (p=1.0000) |
| 3 | structural | -0.039 (p=1.0000) |
| 4 | aec_beta | -0.021 (p=1.0000) |
| 4 | fmri_surface_d25_partial | -0.007 (p=1.0000) |
| 3 | sensor_tde_cov | -0.033 (p=1.0000) |
| 4 | nc_aec_delta | -0.056 (p=1.0000) |
| 2 | nc_aec_theta | -0.103 (p=1.0000) |
| 3 | structural_FS | -0.013 (p=1.0000) |
| 1 | fmri_parcel_partial | 0.012 (p=1.0000) |
| 3 | hmm_covs | 0.004 (p=1.0000) |

C. Proof of the equality between Linear CKA and correlation between fingerprints similarity matrices, with X and Y are feature matrices with shape (n_subjects, n_features);

$$R_{CKA}^{2}=CKA(XX^{T}, YY^{T})=\frac{\left\| Y^{T}X \right\|_{F}^{2}}{\left\| X^{T}X \right\|_{F}\left\| Y^{T}Y \right\|_{F}}$$

$$\frac{\left\| Y^{T}X \right\|_{F}^{2}}{\left\| X^{T}X \right\|_{F}\left\| Y^{T}Y \right\|_{F}}=\frac{tr\left( \left( Y^{T}X \right)^{T}Y^{T}X \right)}{\left\| X^{T}X \right\|_{F}\left\| Y^{T}Y \right\|_{F}}$$

$$=\frac{tr\left( X^{T}YY^{T}X \right)}{\left\| X^{T}X \right\|_{F}\left\| Y^{T}Y \right\|_{F}}=\frac{tr\left( {XX}^{T}{YY}^{T} \right)}{\left\| X^{T}X \right\|_{F}\left\| Y^{T}Y \right\|_{F}}$$

$$=\frac{vec\left( XX^{T} \right)^{T}vec(YY^{T})}{\left\| vec(XX^{T}) \right\|_{2}\left\| vec(YY^{T}) \right\|_{2}}$$

D. results across all features, including no leakage corrected source-space MEG fingerprints:

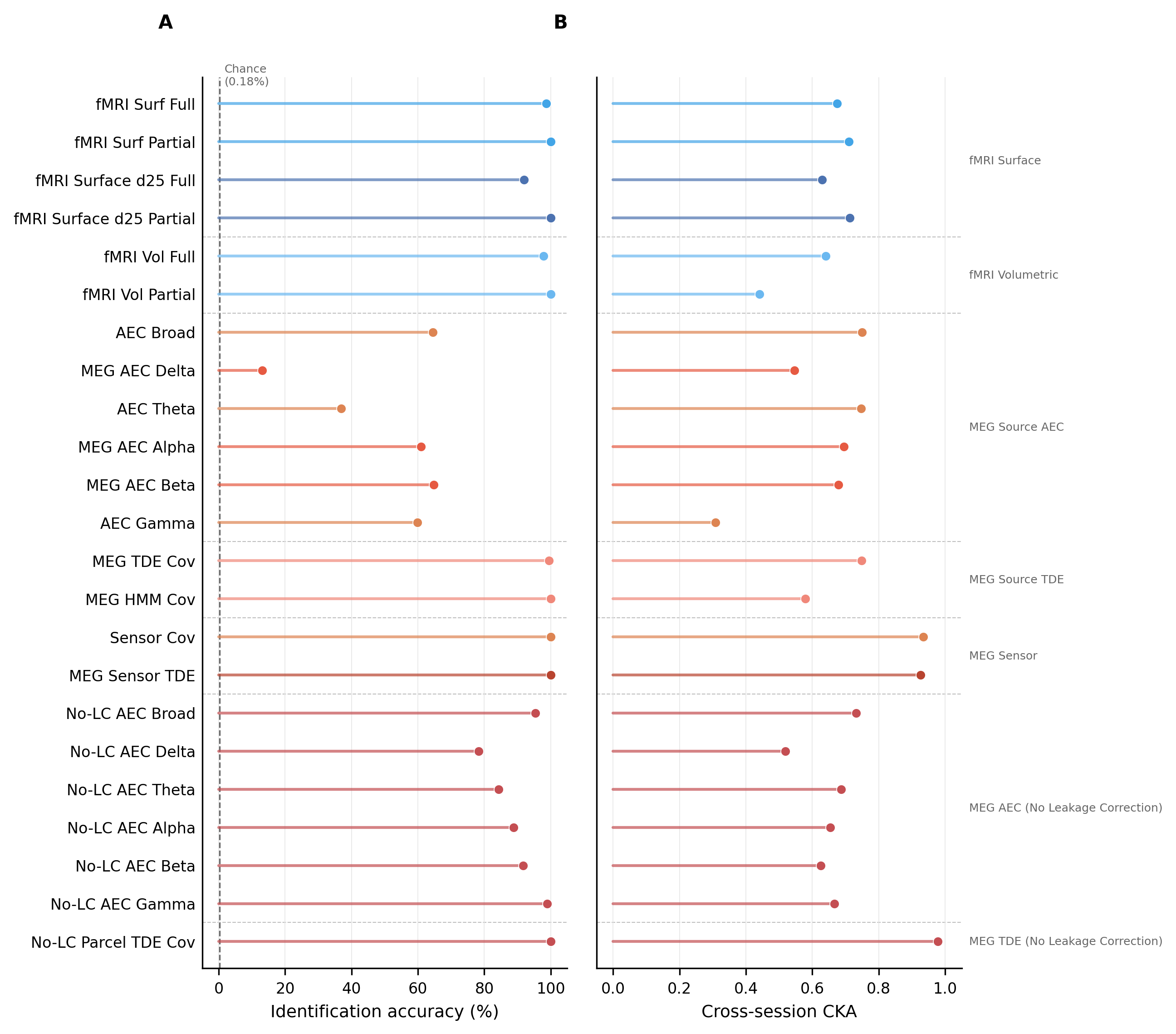

Fig. S1. Cross-session robustness across all features, including no leakage corrected source-space MEG fingerprints.

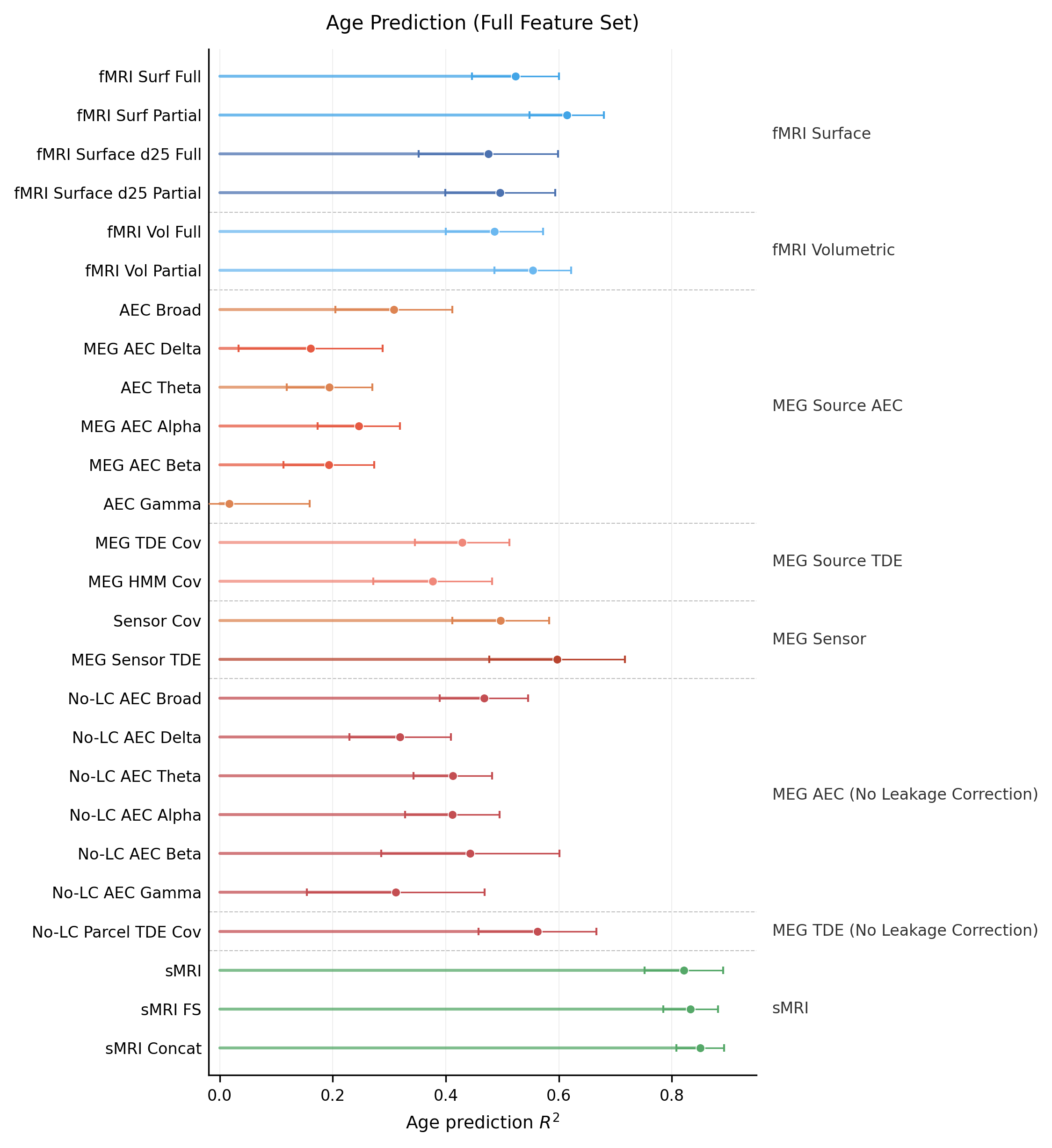

Fig. S2. Age prediction performance across all features, including no leakage corrected source-space MEG fingerprints.

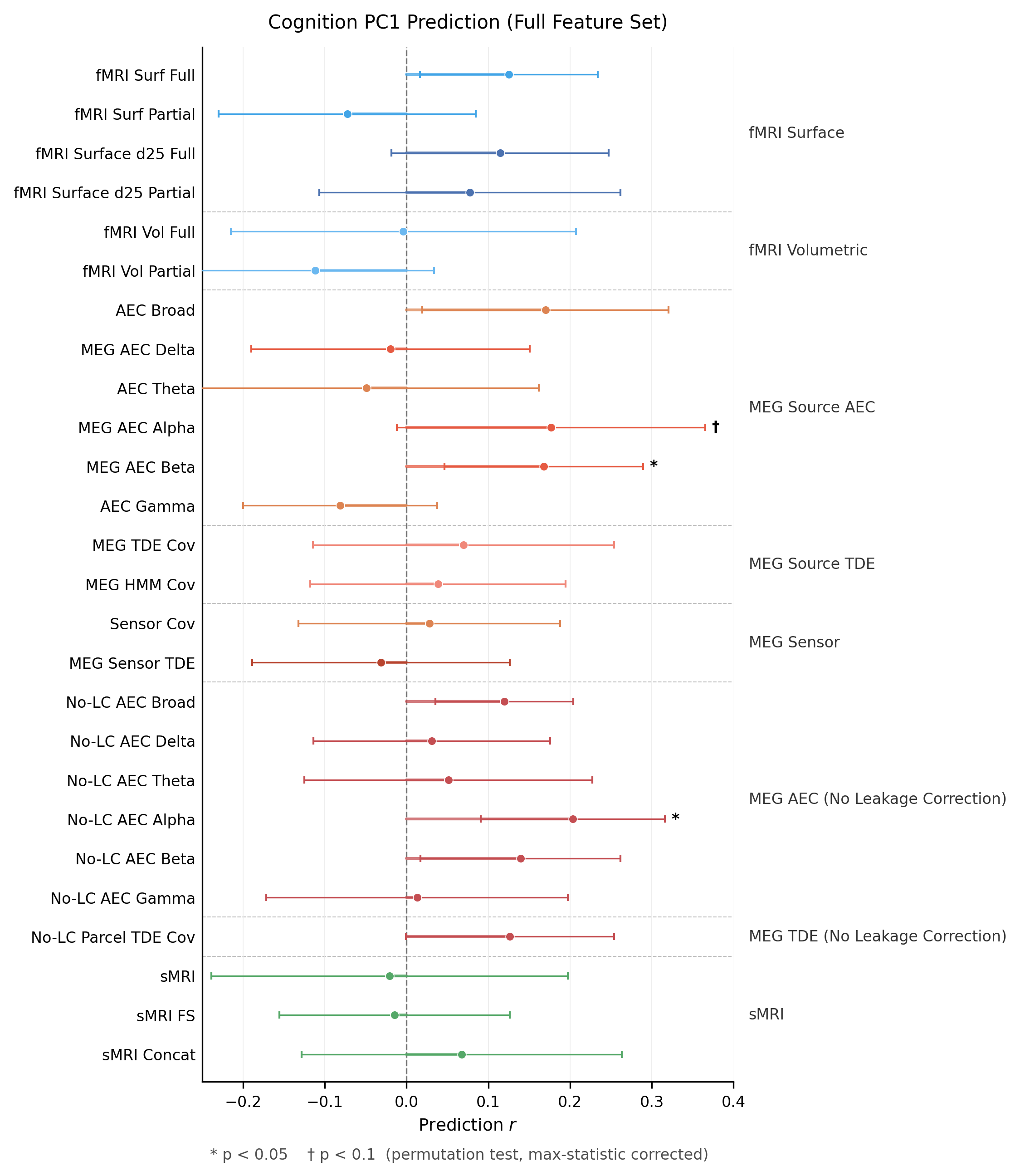

Fig. S3. Cognition PC1 prediction performance across all features, including no leakage corrected source-space MEG fingerprints.

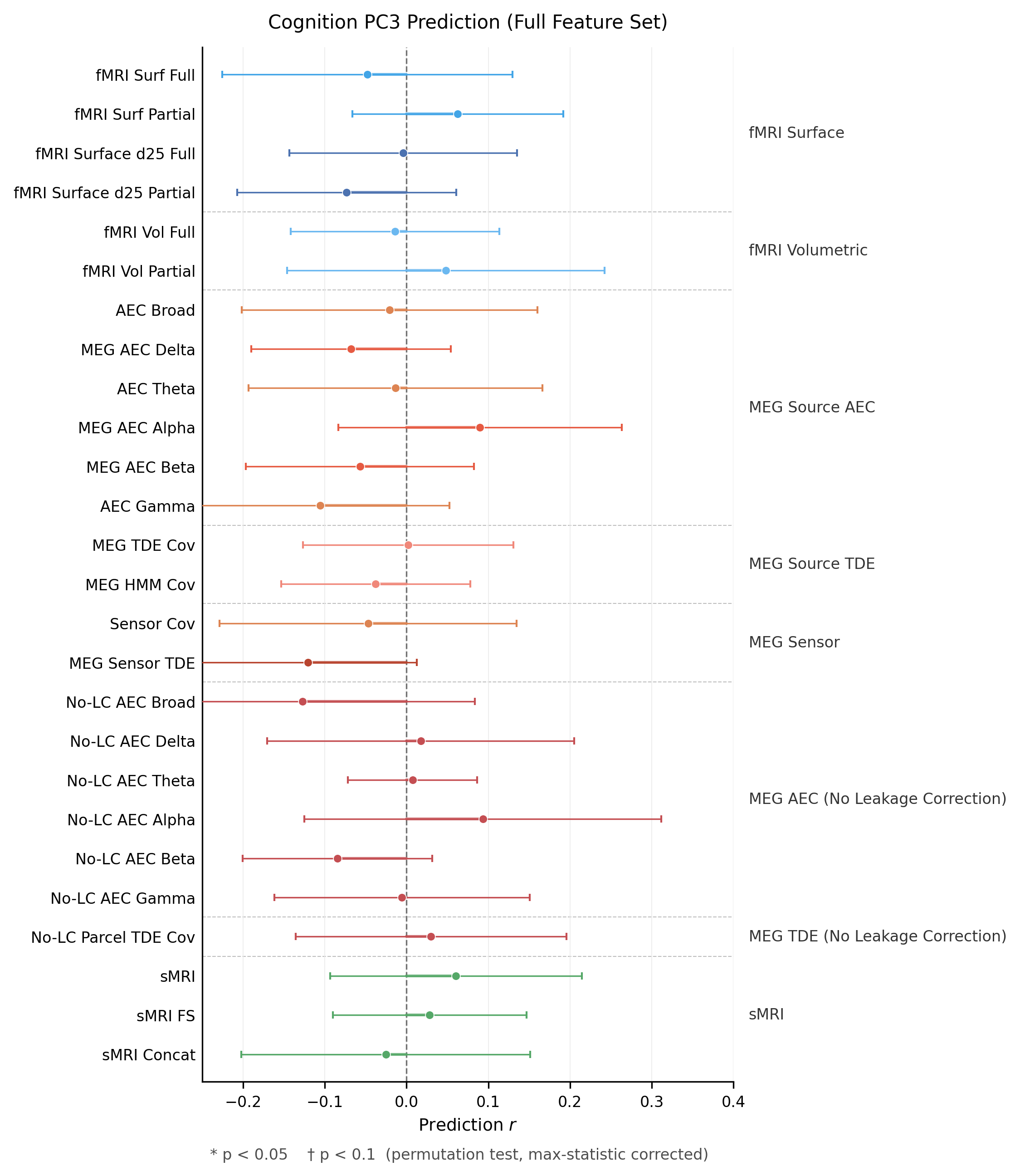

Fig. S4. Cognition PC2 prediction performance across all features, including no leakage corrected source-space MEG fingerprints.

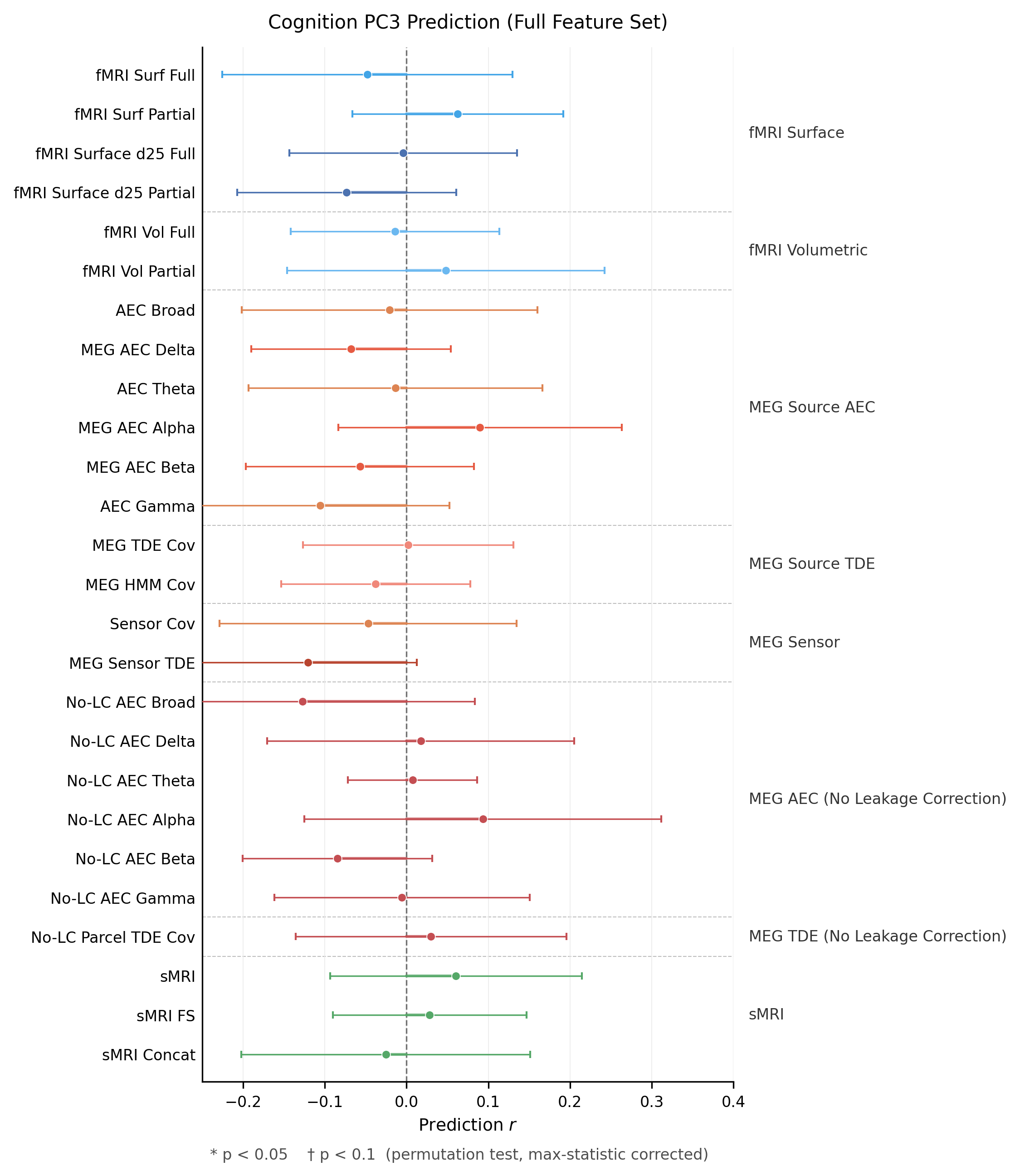

Fig. S5. Cognition PC3 prediction performance across all features, including no leakage corrected source-space MEG fingerprints.

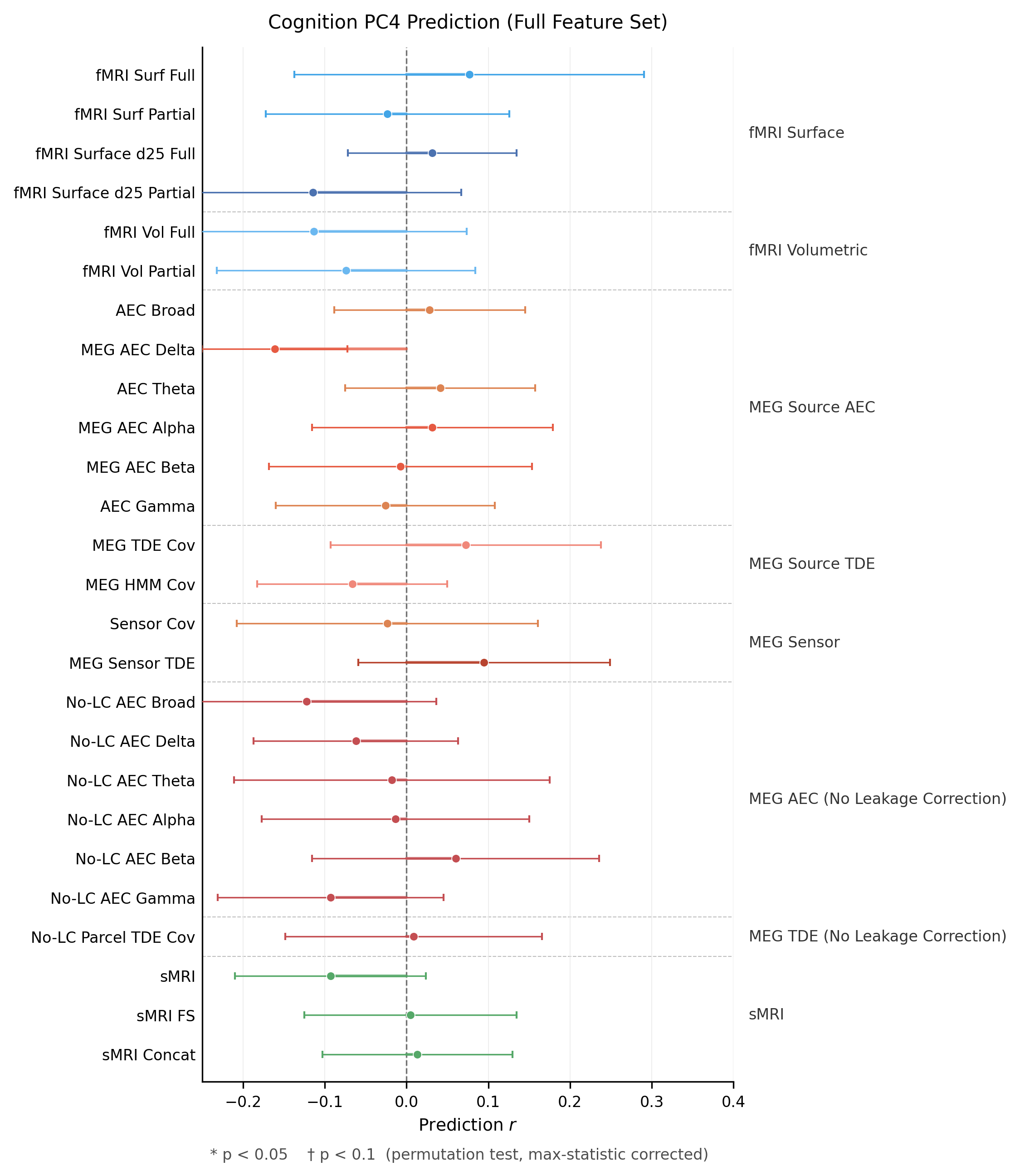

Fig. S6. Cognition PC4 prediction performance across all features, including no leakage corrected source-space MEG fingerprints.

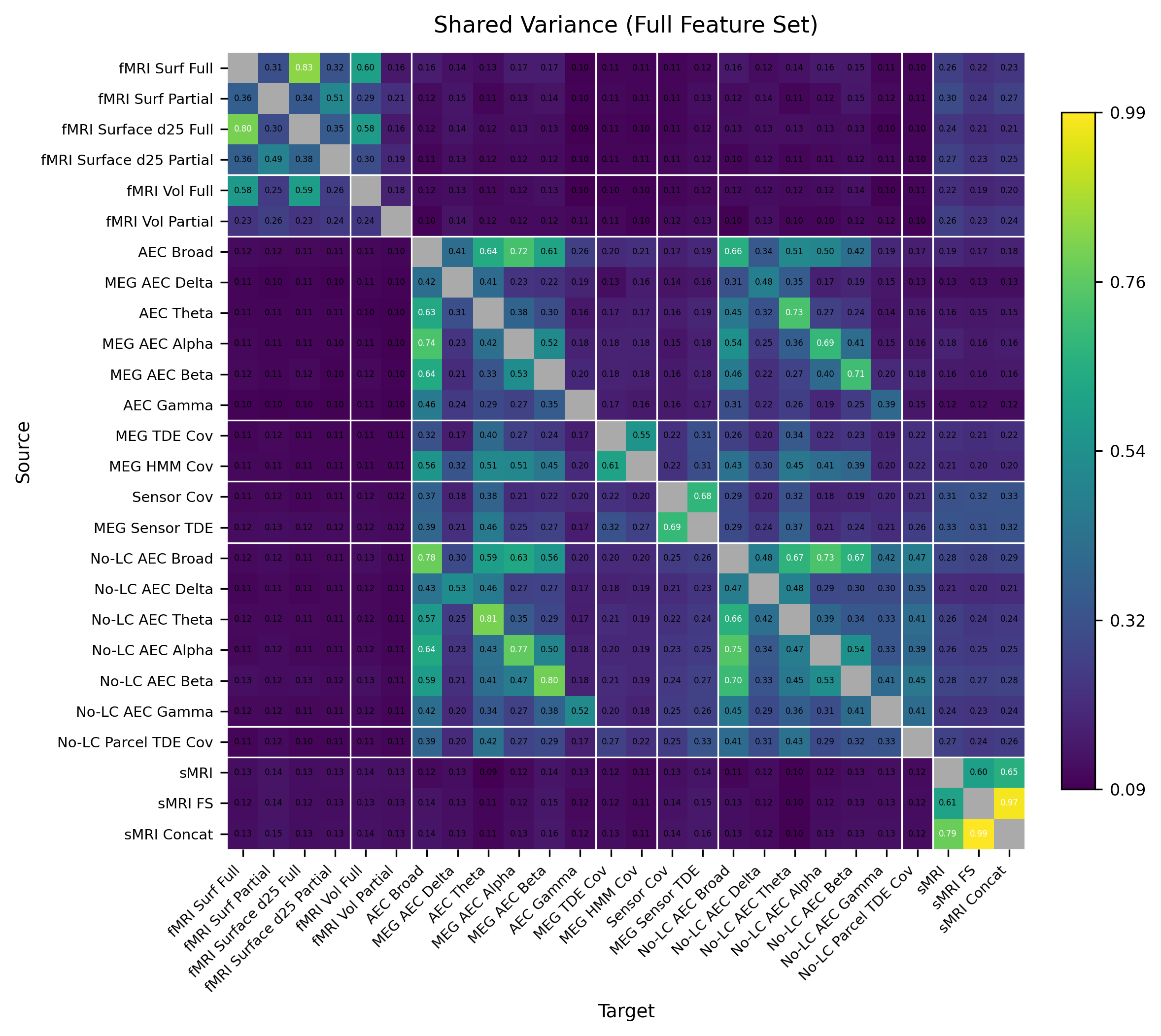

Fig. S7. Full cross-modal shared subspace estimation heatmap across all features, including no leakage corrected source-space MEG fingerprints.

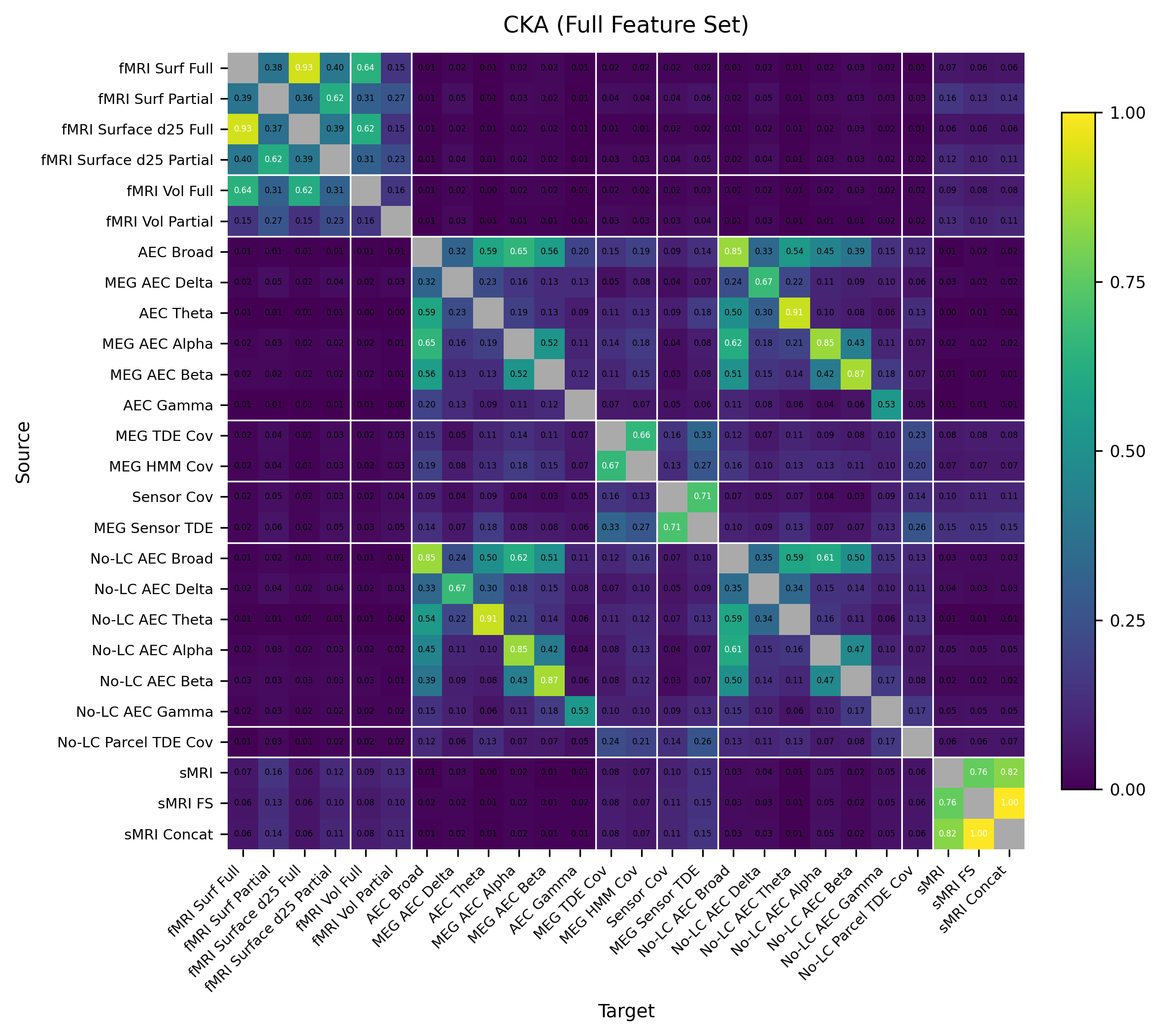

Fig. S8. Full cross-modal CKA results across all features, including no leakage corrected source-space MEG fingerprints.

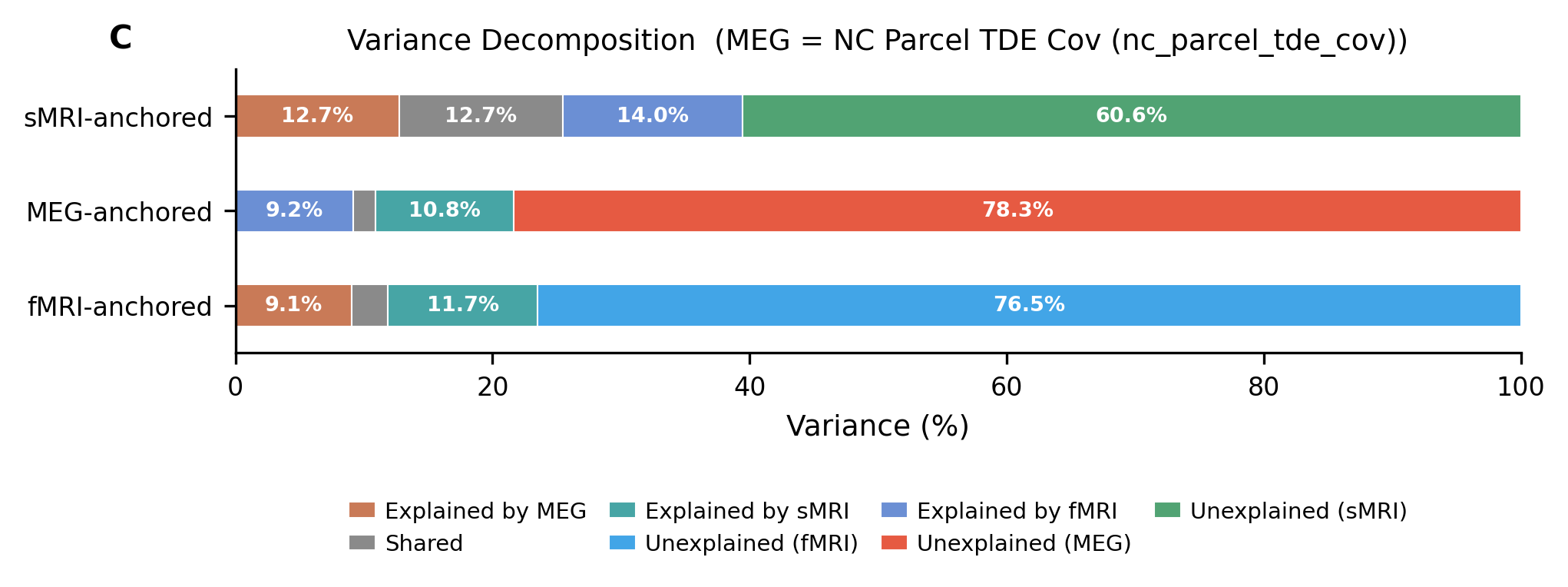

Fig. S9. Directional general variance decomposition with no leakage corrected source space TDE covariances as anchored MEG feature.

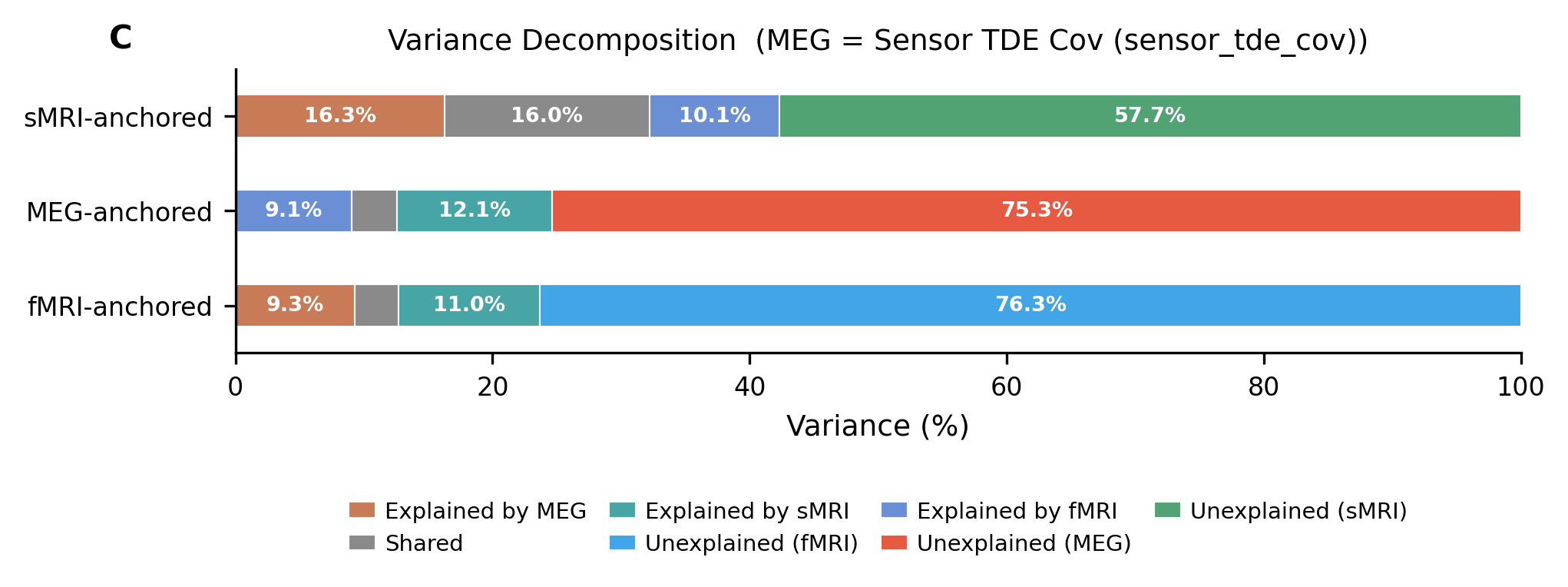

Fig. S10. Directional general variance decomposition with sensor space TDE covariances as anchored MEG feature.

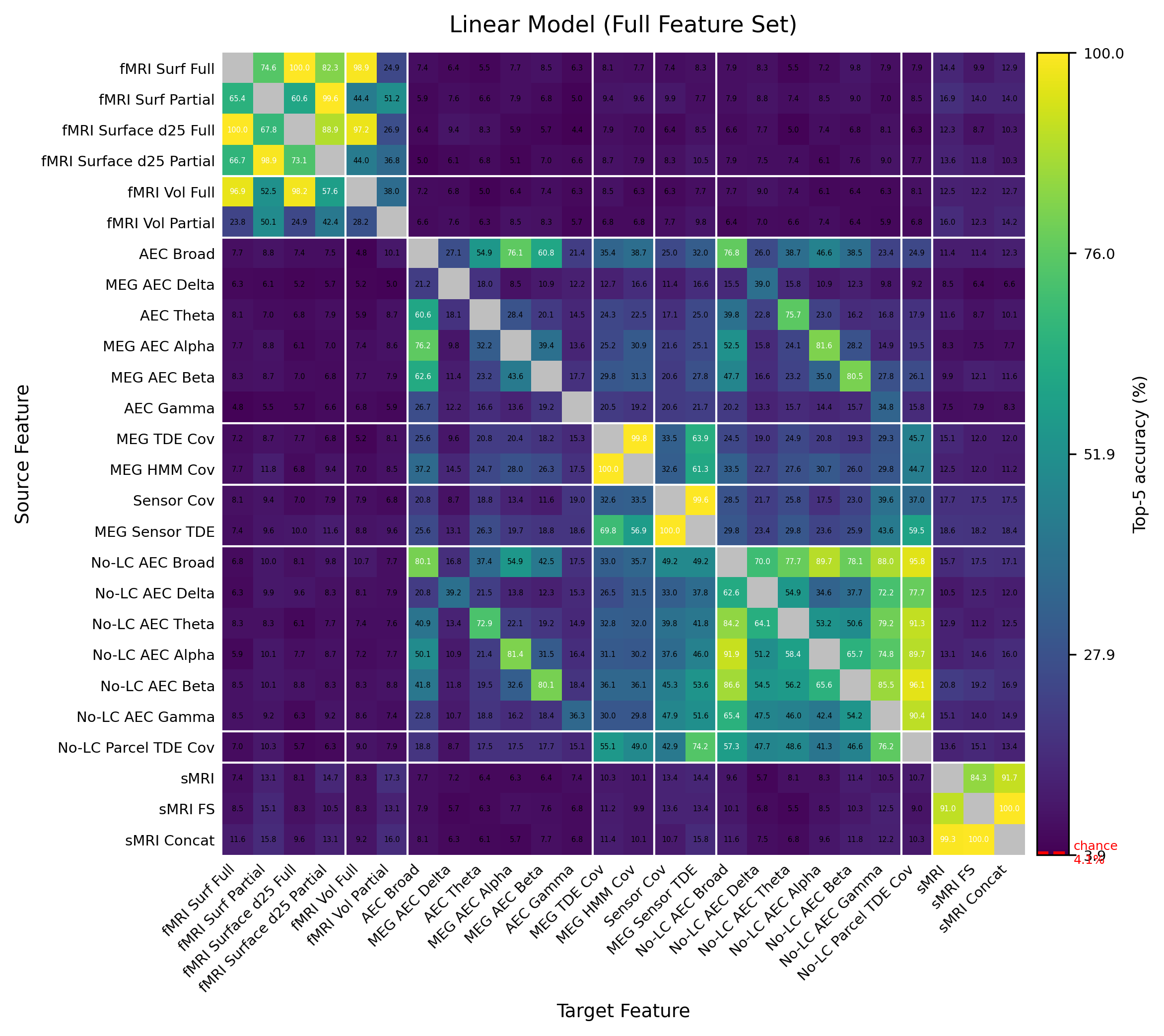

Fig. S11. Full cross-modal subject prediction result with linear model

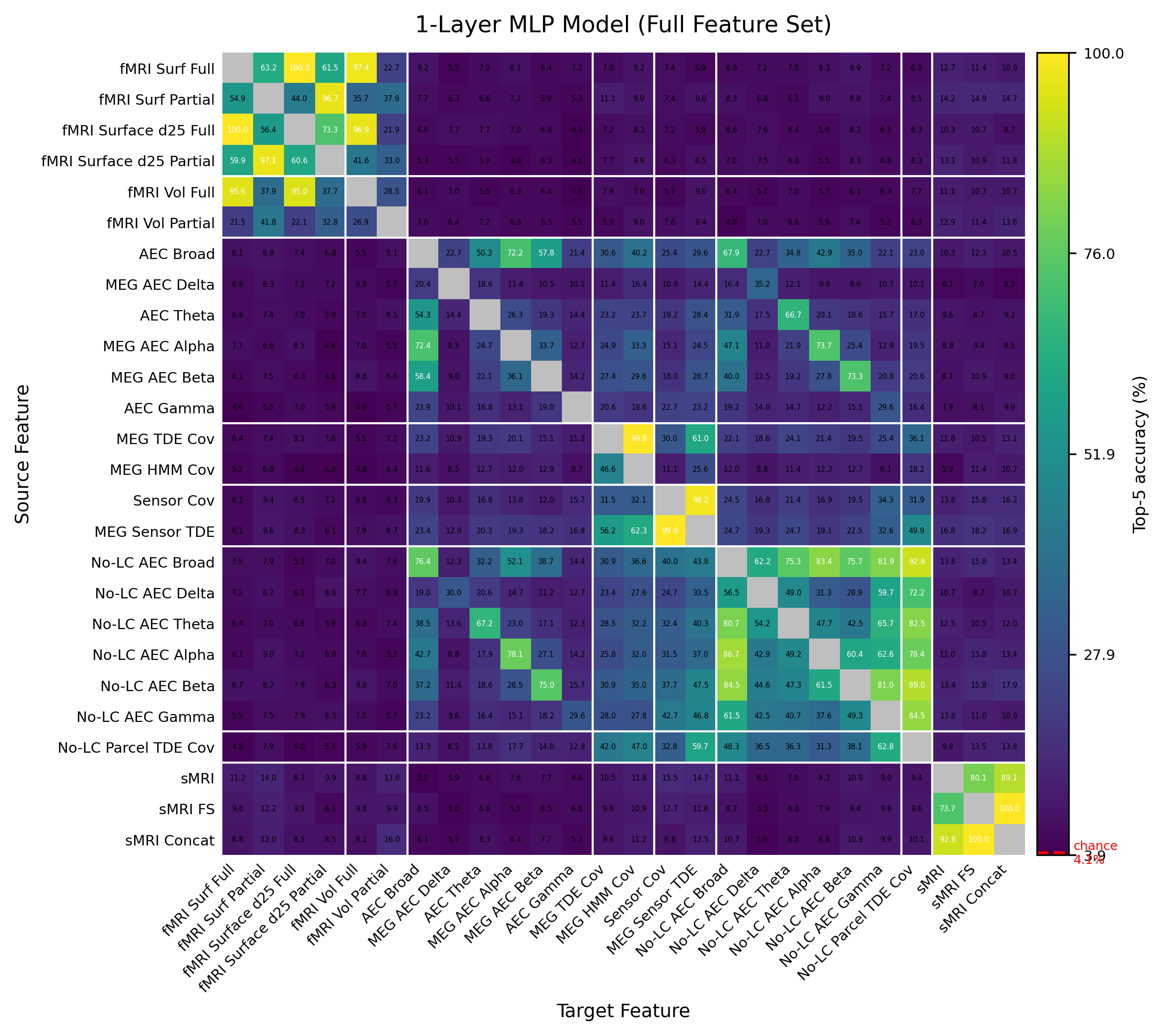

Fig. S12. Full cross-modal subject prediction result with non-linear model
